# UPF1 mutants with intact ATPase but deficient helicase activities promote efficient nonsense-mediated mRNA decay

**DOI:** 10.1101/2022.07.08.499162

**Authors:** Joseph H. Chapman, Jonathan M. Craig, Clara D. Wang, Jens H. Gundlach, Keir C. Neuman, J. Robert Hogg

## Abstract

The conserved RNA helicase UPF1 coordinates nonsense-mediated mRNA decay (NMD) by engaging with mRNAs, RNA decay machinery, and the terminating ribosome. UPF1 ATPase activity is necessary for mRNA target discrimination and completion of decay, but the mechanisms through which UPF1 enzymatic activities such as helicase, translocase, RNP remodeling, and ATPase-stimulated dissociation influence NMD remain poorly defined. Using high-throughput biochemical assays to quantify UPF1 enzymatic activities, we show that UPF1 is only moderately processive (< 200 nt) in physiological contexts and undergoes ATPase-stimulated dissociation from RNA. We combine an in silico screen with these assays to identify and characterize known and novel UPF1 mutants with altered helicase, ATPase, and RNA binding properties. We find that UPF1 mutants with substantially impaired processivity, slower unwinding rate, and reduced mechanochemical coupling (i.e. the ability to harness ATP hydrolysis for work) still support efficient NMD in human cells. These data are consistent with a central role for UPF1 ATPase activity in driving cycles of RNA binding and dissociation to ensure accurate NMD target selection.

## Introduction

The nonsense-mediated mRNA decay (NMD) pathway is a translation-dependent mechanism responsible for both quality control of mRNAs with premature termination codons (PTCs) and general gene expression regulation^1,2^. The RNA helicase UPF1 is the lynchpin of NMD, serving as a protein scaffold for decay complex assembly^3^. Several components of the NMD pathway directly interact with UPF1 to promote decay, including UPF2^4–6^, the PI3K-like kinase SMG1^7,8^, the SMG5/SMG7 heterodimer^9–11^, the endonuclease SMG6^9–13^, and the terminating ribosome^14,15^. In metazoans, phosphorylation of RNA-bound UPF1 by SMG1 is an important pro-decay signaling event, as it promotes assembly of decay enzymes on the mRNA^11–13,16^. These observations suggest that prolonged association of UPF1 with an mRNA is a prerequisite for NMD, as it permits phosphorylation by SMG1, sensing of translation termination, and assembly of RNases on the mRNA.

Mutations in UPF1 that disrupt ATP binding or hydrolysis enable more stable UPF1-RNA binding in the presence of ATP^17,18^ and result in hyperphosphorylated UPF1 in mammalian cells^19^. These features might be expected to enhance NMD, but ATPase-deficient mutants strongly impair NMD in yeast^14,15,17,20,21^ and human^22,23^ cells, suggesting that UPF1 ATPase activity is important for one or more steps in NMD after UPF1 associates with the mRNA. As a superfamily 1 (SF1) helicase, UPF1 uses the energy from ATP hydrolysis to generate work in the form of 5′-3′ translocation on nucleic acids^18,24–27^, duplex unwinding, and disruption of RNA-protein interactions^28,29^. These activities are proposed to aid in recycling ribosomes at PTCs^14,15^, displace RNA-stabilizing factors prior to decay initiation^28^, and disassemble NMD complexes to allow complete mRNA degradation by XRN1^23,30^, a processive 5′-3′ exonuclease^31–34^. Models to explain the requirement for UPF1 enzymatic activity thus span the entire process of decay, from initial substrate selection^35^ to post-decay messenger ribonucleoprotein (mRNP) disassembly^14,15,23^. However, experimental systems capable of distinguishing among specific UPF1 enzymatic features (including ATPase, helicase, translocase, RNP remodeling, and ATPase-stimulated dissociation) and their roles in decay are lacking, limiting mechanistic understanding of the enzymatic roles of UPF1 in NMD.

Here, we develop an experimental framework for 1) quantitative measurement of UPF1 enzymatic properties in vitro and 2) evaluation of the NMD functionality of UPF1 mutants with alterations in those enzymatic properties in human cells. First, using ensemble and single-molecule approaches, we find that UPF1 has only moderate processivity (10s-100s of nucleotides [nt]) when the nucleic acid substrate is not subjected to external forces that destabilize the duplex (e.g. as in magnetic tweezers experiments). We have developed an array of novel high-throughput biochemical assays to quantify the enzymatic properties of UPF1 and computationally identified UPF1 mutants with altered RNA binding properties. Combining these approaches, we quantify the enzymatic properties of a panel of UPF1 ATPase/helicase mutants. We further find that UPF1 mutants with poor processivity, slow unwinding rate, impaired protein displacement activity, and impaired coupling between ATPase activity and unwinding/translocation, efficiently restore NMD upon depletion of endogenous UPF1 in human cells. These data are consistent with a role for UPF1 ATPase activity in sampling potential substrates^35^, but call into question whether efficient mechanochemical coupling and processive RNP remodeling of UPF1 are necessary to conduct NMD.

## Results

### In vitro unwinding assays reveal UPF1 processivity of less than 200 nucleotides (nt) in the absence of destabilizing forces

To better understand how the enzymatic activities of UPF1 impact NMD (**Fig. 1a**), we first modified a recently reported real-time fluorescence-based unwinding assay^29,36^ to measure UPF1 processivity and unwinding rates. We used the helicase core of UPF1 (UPF1-HD) to avoid auto-inhibition by its N-terminal CH^6,37,38^ and C-terminal SQ^39^ domains (**Fig. S1a**). In this assay, UPF1-HD is pre-bound to the fluorescent substrate, and upon addition of ATP, UPF1 either 1) completely unwinds and displaces the fluorescent strand, which then hybridizes to a complementary quencher strand, decreasing the fluorescence or 2) dissociates before complete unwinding (**Fig. 1b**, **Fig. S1b-c**). A lower fluorescence value at the end of the reaction therefore indicates more complete unwinding events. As controls, we observed no unwinding activity with ATPase-deficient DE636AA and K498A ^17,18,21^ UPF1-HD mutants (**Fig. S1d-e**).

**Figure 1.**
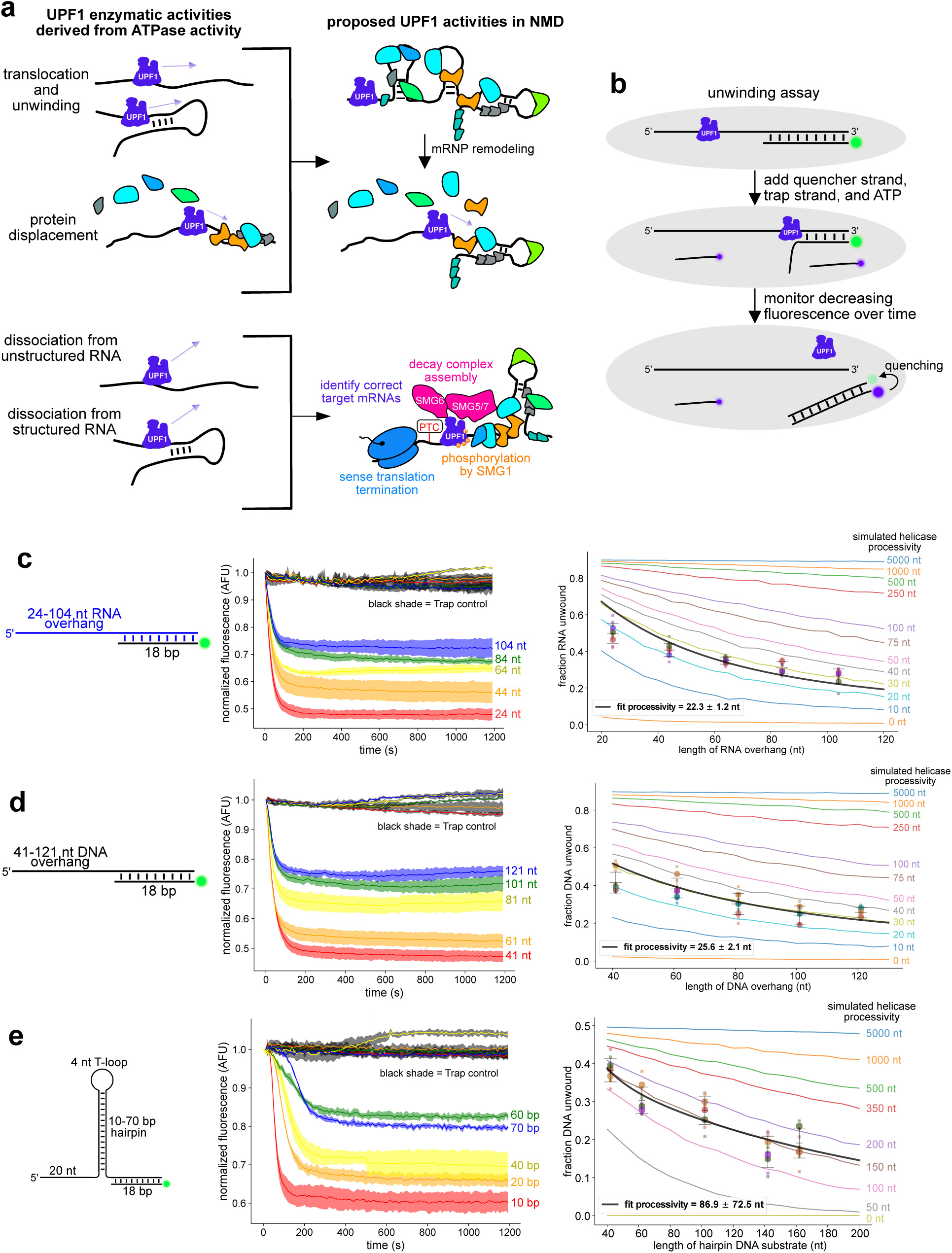
UPF1-HD processivity is less than 200 nt on RNA and DNA. (a) Schematic of UPF1 enzymatic activities on RNA and how they might translate into cellular activities. (b) Single turnover unwinding assay. Trap strand was omitted in multiple turnover assays. (c) Representative time-dependent fluorescence curves from single turnover unwinding measurements with varying lengths of 5′ ssRNA overhang (left). Trap control reactions included 1000-fold excess trap strand prior to UPF1-HD addition. Shaded areas represent standard deviation. Fraction of substrate unwound as a function of the 5′ ssRNA overhang length (right). Each colored dot represents a separate experiment from 3 independent experiments. Small dots represent technical replicates. Black lines indicate processivity model fits and colored lines represent simulations of helicases with differing processivities (see Methods). (d-e) As in (c) but with varying lengths of 5′ ssDNA overhang (d) or hpDNA stem (e).

To account for different nucleic acid structures, we measured UPF1-HD translocation processivities on ssRNA and ssDNA, and unwinding processivities on dsRNA and hairpin DNA (hpDNA). In all cases, we found that UPF1-HD unwound a smaller proportion of substrate as the substrate length increased (**Fig. 1c-e**, **Fig. S1f**, left graphs). We fit the data with established models (**Methods**)^40^ to obtain the processivity for each type of substrate (**Fig. 1c-e**, **Fig. S1f**, black lines on right graphs). We also compared the data to simulated helicases with differing processivities (**Methods**, **Fig. 1c-e**, **Fig. S1f**, colored lines on right graphs). From these analyses, we determined UPF1-HD processivity to be 20-50 nt on ssRNA, ssDNA, and dsRNA and 100-200 nt on hpDNA.

Surprisingly, these processivities are 2-3 orders of magnitude lower than those previously measured using magnetic tweezers, in which hairpin substrates were subjected to duplex-destabilizing forces^28,41,42^. To investigate reasons for this discrepancy, we first performed additional controls to validate the fluorescence-based unwinding assay for processivity measurements. We first corroborated hpDNA processivity by performing an alternate processivity measurement based on a titration of UPF1-HD versus substrate concentration (**Methods**, **Fig. S1g**), and ensured that unwinding was primarily initiated at single-stranded regions (**Fig. S2a**). We next compared the pH 6.0 reaction buffer used in our experiments with the magnetic tweezers pH 7.5 reaction buffer under single (**Fig. 1e**, left graph; **Fig. S2b**) and multiple turnover (**Fig. S2c**) unwinding conditions. This revealed substantially reduced unwinding activity at pH 7.5 compared to pH 6.0, consistent with the acidic preference of human UPF1 in vitro^27^. We further verified that the concentration of the trap strand did not affect unwinding activity (**Fig. S2d**). Lastly, we altered the annealing ratios of substrate strand to fluorescent strand from 11:7 to 1:1 to ensure that the decrease in the fraction unwound on longer substrates was not due to longer unannealed strands acting as more effective UPF1-HD binding sinks (**Fig. S2e-f**). These control experiments reinforced our finding of significantly lower processivity of UPF1-HD in the absence of duplex-destabilizing force.

We next considered the possibility that, despite similar unwinding rates (**Fig. S3a**)^28,42,43^, the UPF1-HD recombinant protein preparation used here was less processive compared to that used in magnetic tweezers experiments. Therefore, we performed unwinding assays on a 535 bp hpDNA under 7-9 pN of force using magnetic tweezers (**Fig. S3b**). In agreement with previous studies^28,41–43^, we observed complete unwinding and translocation through the entire hairpin, indicating a processivity on the order of thousands of nt (**Fig. S3c**). Conversely, without duplex-destabilizing force, UPF1-HD unwound a fluorescent 70 bp hpDNA substrate tethered to a glass surface to a similar extent as in the fluorescence-based unwinding measurements (**Fig. S3d-e**). This suggests that the applied duplex-destabilizing force in magnetic tweezers, not tethering the substrate to the surface, is responsible for higher observed UPF1-HD processivity in magnetic tweezers. These observations are consistent with similar observations of force-dependent processivity enhancement of several RNA and DNA helicases ^44–49^.

Lastly, we investigated whether applying force on the nucleic acids in a non-duplex-destabilizing orientation would significantly increase UPF1-HD processivity. Therefore, we performed Single-molecule Picometer Resolution Nanopore Tweezers (SPRNT) unwinding and translocation measurements^50–52^. In SPRNT, a single MspA nanopore in a phospholipid bilayer separates two electrolyte solutions. When a voltage is applied to the system, current flows through the nanopore, drawing the negatively charged DNA pre-bound to UPF1-HD into the nanopore. UPF1-HD controls the passage of the DNA through the nanopore, allowing measurement of UPF1-HD unwinding and translocation via the sequence-dependent changes in current caused by DNA moving through the nanopore (**Fig. 2a-b**). Unlike magnetic tweezers, the applied force in SPRNT does not destabilize the duplex but instead assists UPF1-HD translocation by pulling the DNA through the nanopore.

**Figure 2.**
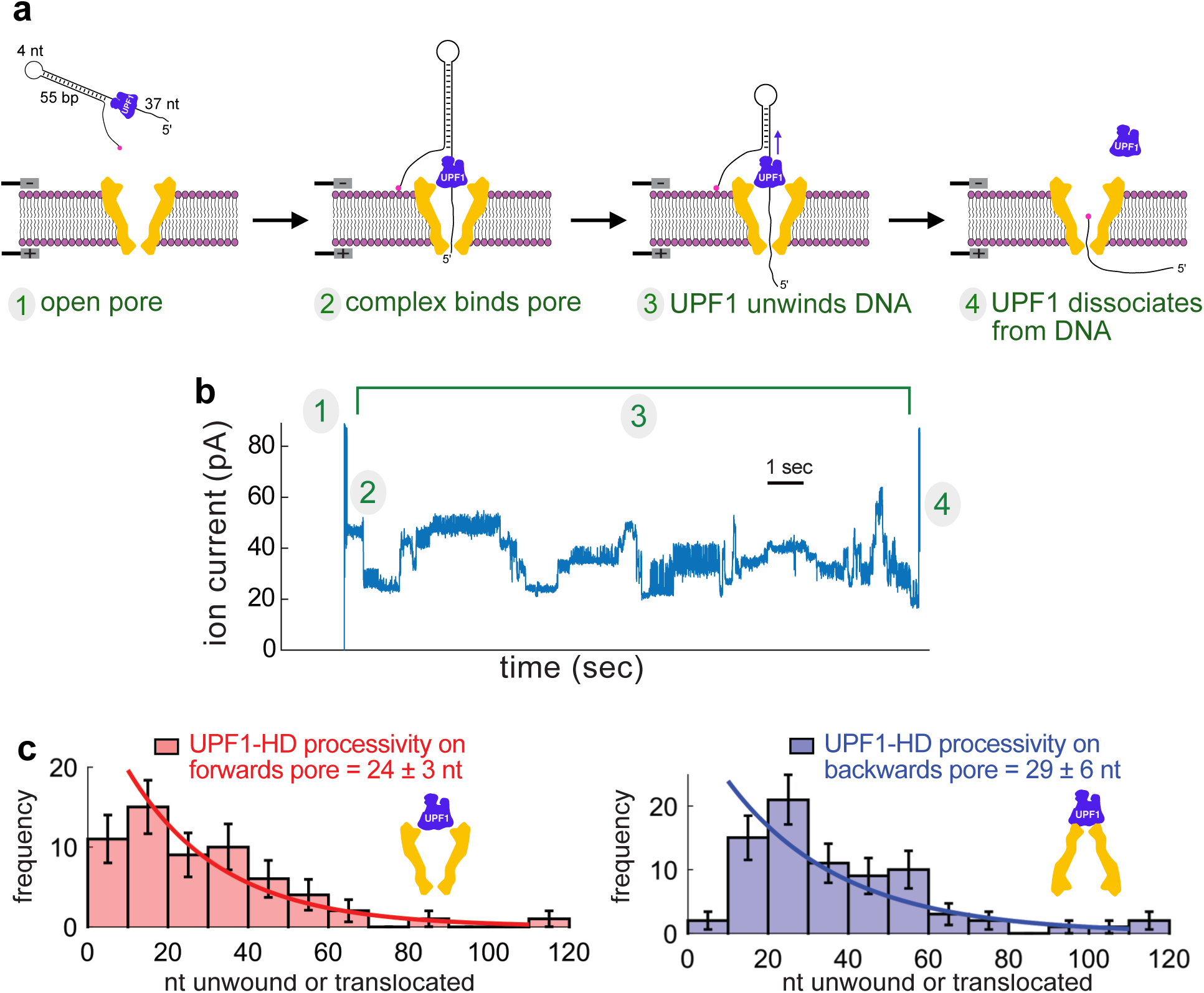
UPF1-HD exhibits low processivity in a single-molecule picometer-resolution nanopore tweezers (SPRNT) system. (a) Schematic of SPRNT system with a “forwards” pore setup. In “backwards” pore experiments, the MspA nanopore (yellow) is inverted 180° relative to the phospholipid bilayer. (b) Example unwinding event indicating the time-dependent ion current associated with the four stages of SPRNT signal illustrated in (a). The open pore current is high (1). The current drops when a UPF1-HD/DNA complex enters the pore (2). The current levels subsequently fluctuate as UPF1 unwinds the hairpin and the 5′ end of the DNA translocates through the pore (3). The current level rapidly rises when UPF1 dissociates and the DNA is pulled unimpeded through the pore (4). (c) Histogram of unwinding/translocation events and processivity fits for forwards (red) and backwards (blue) pore.

We used changes in current to temporally identify UPF1-HD-bound DNA initially docking on the nanopore, UPF1 unwinding and translocation, and subsequent UPF1-HD dissociation (**Fig. 2a-b**). We aligned experimental traces (**Fig. S4a**) to the predicted current (**Fig. S4b**) to determine the position of UPF1-HD over time (**Fig. S4c**) and observed a processivity of ∼24 nt on hpDNA across multiple applied forces (**Fig. 2c**, red). Since UPF1-HD interaction with the MspA nanopore may influence activity, we inverted MspA 180° relative to the membrane and observed similar processivity of ∼29 nt (**Fig. 2c**, blue; **Fig. S4d**), but significantly higher unwinding and translocation rates, which increased even further with increasing assisting force (**Fig. S4e-f**). These results are consistent with fluorescence-based unwinding measurements of UPF1-HD processivity, and suggest that the non-duplex-destabilizing force in SPRNT does not enhance UPF1-HD processivity.

Taken together, our data suggest that UPF1 acts locally, associating and dissociating from cellular RNPs. Our findings are consistent with the hypothesis that the high processivity of UPF1-HD in magnetic tweezers experiments is due to force-induced processivity enhancement, as observed with other SF1^44–47^, SF2^49^, and SF4^48^ helicases. Nevertheless, our results, together with the ability of UPF1 to generate sufficient force to disrupt biotin-streptavidin interactions^28,29^, are compatible with a role for local mRNP remodeling by UPF1 for efficient NMD.

### UPF1 undergoes frequent ATPase-stimulated dissociation events

Since the duration of UPF1 association with mRNA (residence time) has been hypothesized to be an important determinant of NMD specificity and efficiency^35^, we next set out to measure UPF1-HD dissociation from nucleic acids. Although it is possible to estimate residence time from processivities and unwinding rates extracted from fluorescence-based unwinding assays, incomplete unwinding events may confound the results. Therefore, we developed a fluorescence <u>a</u>nisotropy-based <u>d</u>issociation (FAD) assay to monitor UPF1-HD dissociation from fluorescent nucleic acids over time (**Fig. 3a**). In this assay, each fluorescent substrate is pre-bound with ∼5 UPF1-HD molecules (**Fig. S5a**) to produce initial high fluorescence polarization, which subsequently decreases as UPF1-HD dissociates. UPF1-HD remained stably bound in the absence of ATP but underwent ATP-stimulated dissociation, with a half-life of ∼1 min (**Fig. 3b-c**, **S5b**). In control measurements, ATPase-deficient DE636AA and K498A both remained stably bound in the presence and absence of ATP (**Fig. S5c**). If UPF1-HD processivity on ssDNA was high, a large population would translocate to the end of substrates before dissociating, and slower dissociation would be observed with longer 3′ overhangs. However, we did not see significant differences in ATP-stimulated dissociation from substrates with 3′ overhangs over 40 nt (**Fig. 3d**), confirming low processivity and indicating that the assay predominantly monitors UPF1-HD dissociation from internal sites.

**Figure 3.**
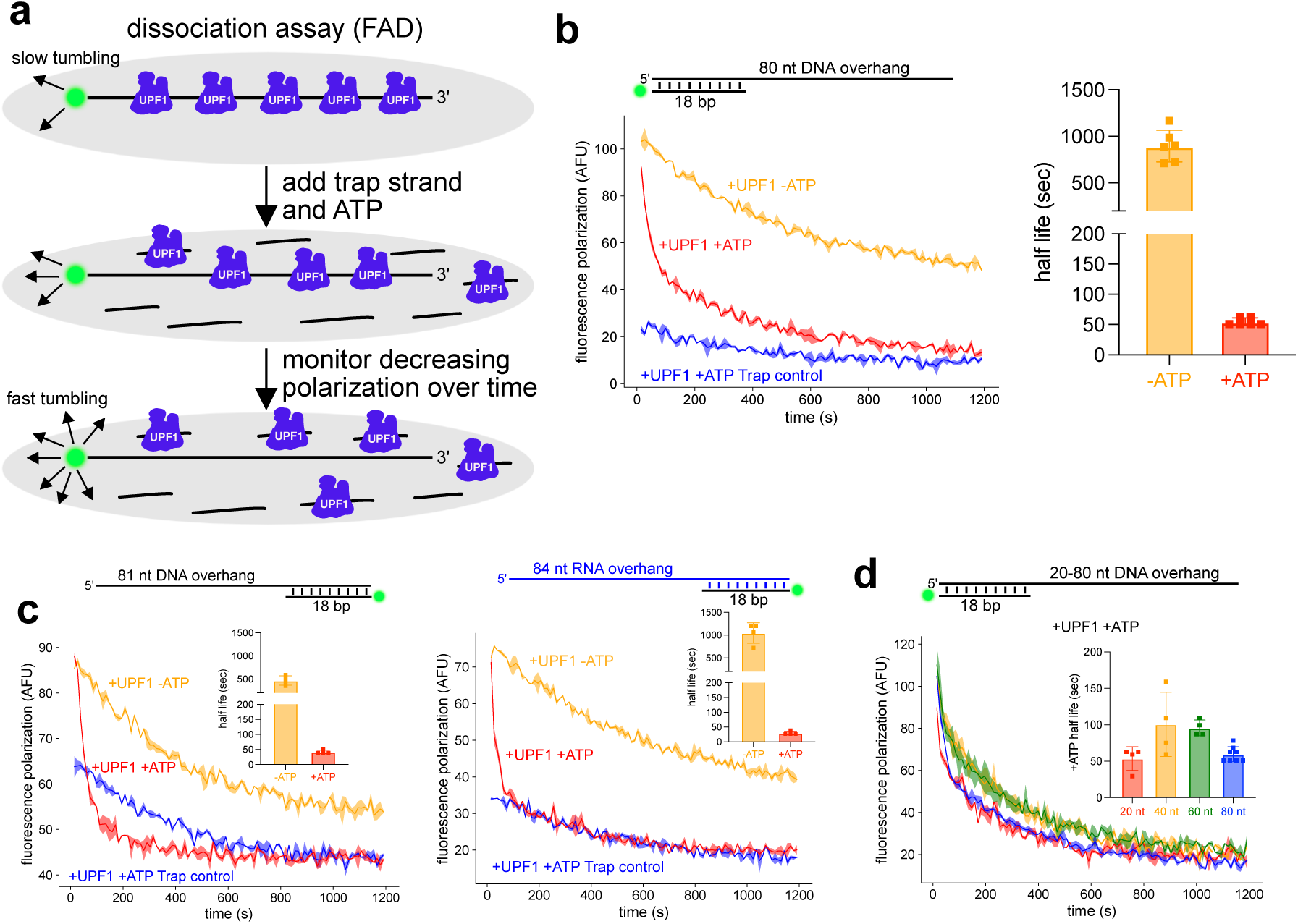
UPF1-HD undergoes ATP-stimulated dissociation from DNA and RNA. (a) Schematic of fluorescence <u>a</u>nisotropy-based <u>d</u>issociation (FAD) assay. More arrows on fluorophore indicates faster tumbling and thus lower polarization. (b) Representative curves from FAD with a 3′ overhang substrate from 4 independent experiments (left). Trap control reactions included 1000-fold excess trap strand prior to UPF1-HD addition. Quantification of binding half-life (see Methods) from the time-dependent fluorescence polarization curves (right). (c) As in (b), but with a 5′ DNA (left) or RNA (right) overhang from 2 independent experiments. Insets indicate half-life quantification. (d) Representative curves from FAD with four different 3′ overhang substrates from 2 independent experiments. For clarity, only +ATP curves are shown. Inset indicates half-life quantification.

Since cellular RNAs are structured, we next monitored UPF1-HD dissociation from structured substrates. Although the greater mass of hairpin substrates limited the dynamic range of the assay, we observed ATP-stimulated dissociation (**Fig. S5d**). As with substrates containing 3’ ssDNA overhangs, UPF1-HD dissociation rates from hairpin substrates plateaued at longer substrate lengths (**Fig S5e**). As an orthogonal approach, we developed a <u>s</u>imultaneous <u>u</u>nwinding and <u>d</u>issociation (SUD) assay by modifying the unwinding assay to include fluorescent UPF1-HD (UPF1-HD-647) and a second quencher strand to capture and reduce the fluorescence of UPF1-HD-647 after dissociation from the substrate (**Fig. S5f**). This assay recapitulated ATP-stimulated dissociation of UPF1-HD, with comparable kinetics (**Fig. S5g**). Consistent with a processivity < 200 nt on hpDNA and the results of the FAD assay (**Fig. S5e**), UPF1-HD dissociation rates plateaued at longer hairpin lengths (**Fig. S5h**). Taken together, these assays allow direct measurement of UPF1-HD dissociation from nucleic acids and indirectly confirm processivity < 200 nt.

### UPF1 ATPase activity is tightly coupled to unwinding and translocation, and dissociation is driven primarily by ATP hydrolysis

To begin to dissect relationships among UPF1 RNA binding, ATPase activity, and mechanochemical coupling, we first performed NADH-coupled ATPase assays^29^ (**Fig. S6a-c**). In conjunction with the fluorescence-based unwinding assays described above, ATPase measurements allow quantification of the ability of UPF1 to couple ATP hydrolysis with work in the form of unwinding and translocation. We observed tight coupling of ∼1 ATP hydrolyzed per nucleotide unwound or translocated (**Fig. S6d**). As controls, we used DE636AA and K498A mutants, both of which expectedly lacked detectable ATPase activity (**Fig. S6e-f**). We next quantified nucleic acid binding affinity using fluorescence anisotropy. In these equilibrium binding assays, we obtained an apparent K_D_ of 0.5-2 nM for UPF1-HD in the absence of ATP (**Fig. S6g-h**), representing a binding affinity 1-2 orders of magnitude tighter than previously reported fluorescence anisotropy binding assays^38,42^. To investigate this discrepancy, we considered known factors that limit apparent equilibrium binding affinities, including substrate concentrations that exceed the K_D_ and incubation times that are insufficient to reach equilibrium^53^. Indeed, we found that letting the system reach equilibrium by increasing incubation time (**Fig. S6i-j**) and decreasing substrate concentration (**Fig. S6k-l**), both significantly increased the measured binding affinity of UPF1-HD for nucleic acids.

It is currently unclear whether ATP binding or ATP hydrolysis regulate UPF1-HD association and dissociation from RNA. In agreement with previous equilibrium binding assays^21,37–39^, we found that ATP binding (using the non-hydrolyzable ATP analog, AMP-PNP) reduced UPF1-HD binding affinity for nucleic acids, but not to the same extent as ATP hydrolysis (using ATP, **Fig. S7a-c**). Furthermore, consistent with previous kinetics experiments^16^, only ATP, not AMP-PNP, promoted dissociation of UPF1-HD from substrates in the FAD assay (**Fig. S7d-e**). These data indicate that ATP binding may impair UPF1 association with RNA but that ATP hydrolysis is the primary driver of UPF1 dissociation from RNA.

### In silico Rosetta-Vienna RNP ΔΔG method identifies known and novel mutations affecting UPF1 RNA binding properties

With a quantitative view of UPF1-HD enzymatic activities in vitro, we next set out to discover new UPF1-HD mutants with alterations in enzymatic activities and characterize them alongside previously identified ATPase and helicase mutants using the high-throughput assays we developed. Therefore, we used the Rosetta-Vienna RNP ΔΔG method^54,55^ to computationally identify UPF1-HD point mutants with RNA binding energies significantly different from WT UPF1-HD (**Fig. 4a**). The predicted RNA binding energies matched expectations for previously characterized “grip mutants” found to have significantly lower processivities in magnetic tweezers experiments (A546H/K547P/S548A [AKS-HPA], A546H, K547P, R549S)^41^, indicating the validity of this approach (**Fig. 4b**). Of the 11,818 point mutants tested, approximately 96% had no effect on predicted RNA binding energy (−1 < ΔΔG < 1, in units of kcal/mol), and most predicted large effects trended towards weaker binding (ΔΔG > 1), with only 29 mutations with predicted stronger binding (ΔΔG < -1, **Fig. S8a**, **Table S1**). We characterized several of these mutants in vitro, then selected a subset for cellular characterization, including two novel mutants with -ΔΔG predictions (E797R, G619K), four of the previously characterized grip mutants^41^ (all +ΔΔG predictions), and the R843C dominant-negative mutant^16,56^ (**Fig 4c**).

**Figure 4.**
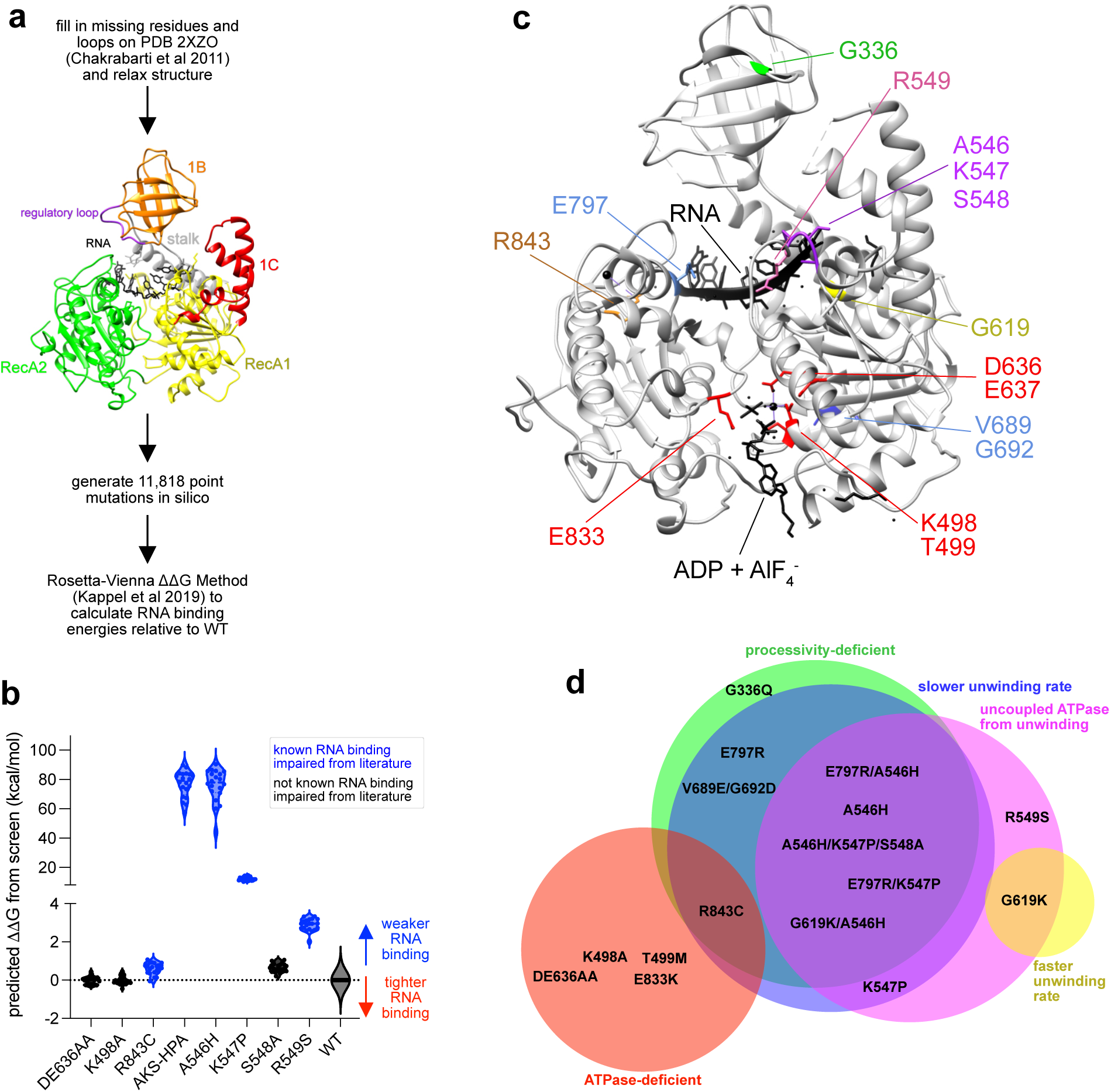
Rosetta-Vienna ΔΔG Method identifies previously characterized grip mutants and novel UPF1-HD mutants with altered RNA binding properties. (a) Workflow of in silico screen to identify UPF1 mutants of interest (see Methods). (b) Change in the binding free energy (ΔΔG) predicted by the screen for previously identified mutants. ΔΔG for each mutant was obtained through comparison against the 20 lowest energy relaxed WT structures. (c) Structure of UPF1-HD indicating selected mutants. Colors correspond to classification in (d). (d) Qualitative Venn diagram generated with DeepVenn^60^ of mutants from in vitro characterization (see Table S2 for quantitative values).

### A546H and G619K/A546H UPF1-HD mutants exhibit severe biochemical defects

After purifying (**Fig S8b-c**) and characterizing these mutants, we identified several overlapping enzymatic classes, including UPF1-HD mutants with altered ATPase rates, processivities, unwinding rates, and ATPase coupling efficiencies (**Fig. 4d**). Aside from the ATPase-deficient mutants, the most severe defects were observed with the previously characterized grip mutant, A546H (AKS-HPA showed similar defects). Consistent with previous reports^41^, A546H exhibited significantly lower processivity and slower unwinding rate (**Fig. S8d**, top; **Table S2**). A546H also showed ∼25-fold less efficient coupling between ATP hydrolysis and unwinding, and modestly lower equilibrium binding affinity (**Fig. S8e**, top; **Table S2**). Additionally, A546H showed faster ATP-independent dissociation as monitored by the FAD assay, but continued to undergo ATP-stimulated dissociation from 3′ ssDNA overhang substrates (**Fig. S8f**, top; **Table S2**). In contrast, A546H did not undergo ATP-stimulated dissociation from substrates with 5′ ssDNA overhangs, even though A546H ATPase activity was enhanced when bound to such structured substrates (**Fig. S8g**, top; **Table S2**). This effect was even more pronounced on substrates with 5′ ssRNA overhangs (**Fig. 5a**), in which ATP simulated dissociation of WT UPF1-HD (**Fig. 5b**, top), but slowed down dissociation of A546H from these substrates (**Fig. 5b**, middle). We reasoned that this could be caused by decreased dissociation of A546H molecules that had translocated into the duplex region. The A546 half-life on the substrate would thus consist of two populations, one undergoing fast dissociation from single-stranded regions and one undergoing slow dissociation from the duplex region. In the absence of ATP, A546H would be unable to access the duplex region, so fast dissociation from the ssRNA would predominate.

**Figure 5.**
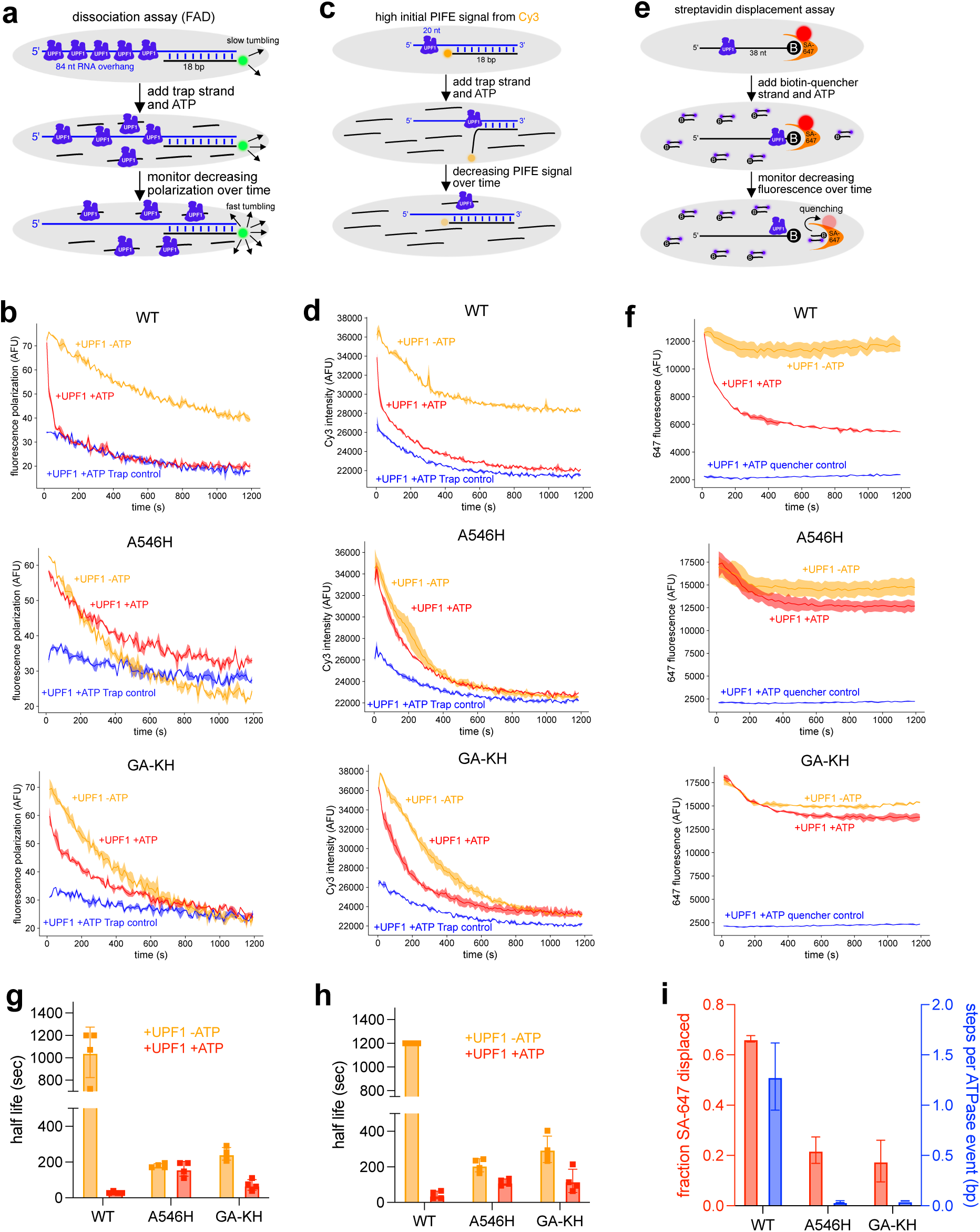
Dissociation and PIFE assays indicate differential release kinetics of UPF1-HD mutant proteins. (a) FAD assay with indicated 5′ RNA overhang substrate. (b) Representative curves from FAD of WT (top), A546H (middle), and GA-KH (bottom) UPF1-HD constructs from 2 independent experiments. (c) Schematic of PIFE assay with indicated 5′ RNA overhang substrate. (d) Representative curves from PIFE assay of WT (top), A546H (middle), and GA-KH (bottom) from 2 independent experiments. Trap control reactions were performed by adding 1000-fold excess trap strand prior to UPF1-HD addition. (e) Fluorescence-based biotin-streptavidin displacement assay. (f) Representative curves of biotin-streptavidin displacement assay of WT (top), A546H (middle), and GA-KH (bottom) from 2 independent experiments. Quencher control reactions were performed by adding 63-fold excess biotin-quencher strand prior to streptavidin addition. (g-h) Half-life quantification from (b) and (d). (i) Fraction streptavidin displaced quantified from (f) and ATPase coupling calculated by dividing unwinding rate by ATPase rate on hpDNA. Error bars indicate standard deviation.

To test this hypothesis, we monitored the presence of UPF1-HD near the single-strand/double-strand junction using protein-induced fluorescence enhancement (PIFE), a phenomenon caused by enhanced fluorescence intensity of dyes such as Cy3 that undergo cis-trans isomerization when in proximity to proteins (**Fig. 5c**)^57^. In control PIFE assays, WT UPF1-HD bound at the junction caused enhanced Cy3 fluorescence intensity, which decreased over time only in the presence of ATP as the protein dissociated from or translocated through the RNA (**Fig. 5d**, top). Consistent with impaired ATP-stimulated dissociation from the duplex region, A546H did not exhibit ATP-stimulated PIFE decrease over time (**Fig. 5d**, middle). Together, the ATPase, FAD, and PIFE assays suggest A546H cannot efficiently release from structured regions, where it instead unproductively hydrolyzes ATP.

Since A546H did not efficiently couple ATP hydrolysis into work in the form of unwinding, we reasoned that this should also result in reduced force generation and, in turn, impaired displacement of proteins from nucleic acids. As a stringent test of protein displacement, we developed a fluorescence-based biotin-streptavidin displacement assay, in which fluorescence decreases as fluorescent streptavidin is displaced from the biotinylated DNA (**Fig. 5e**). Consistent with previous reports^28,29^, WT UPF1-HD efficiently disrupted biotin-streptavidin interactions only in the presence of ATP (**Fig. 5f**, top). Importantly, A546H streptavidin displacement activity was significantly less efficient than that of WT UPF1-HD (**Fig. 5f**, middle), consistent with the idea that it generates less work from each ATPase cycle.

In an attempt to enhance the helicase activity of the A546H mutant, we combined it with the G619K mutation, which exhibited increased rates of ATPase, unwinding, and dissociation, to generate G619K/A546H (GA-KH). Relative to A546H, GA-KH showed modestly enhanced unwinding activity, slightly faster unwinding rate, and similar decoupling of ATPase activity from unwinding (**Fig. S8d-e**, bottom; **Table S2**). Notably, GA-KH exhibited enhanced ATP-stimulated dissociation relative to A546H, as shown by both FAD (**Fig. 5b**, bottom; **Fig. 5g**, **Fig. S8f-g**, bottom) and PIFE (**Fig. 5d**, bottom; **Fig. 5h**) assays. Consistent with decoupled ATPase activity from unwinding, GA-KH exhibited similar streptavidin displacement activity compared to A546H (**Fig. 5f**, bottom; **Fig. 5i**). Taken together, both A546H and GA-KH exhibit similar biochemical defects, although GA-KH has modestly higher unwinding activity and restores ATP-simulated dissociation.

### Slow, poorly processive UPF1 mutants with mechanochemical decoupling support efficient NMD in human cells

After quantifying the biochemical aspects of A546H and GA-KH in addition to other UPF1-HD mutants (**Fig. 4d**, **Fig. S9a-d**, **Table S2**), we set out to determine their NMD efficiency in human cells. To this end, we generated stable cell lines expressing siRNA-resistant CLIP-tagged UPF1 (CLIP-UPF1) mutants in Flp-In T-Rex 293 cells, with or without siRNA-induced endogenous UPF1 knockdown (**Fig 6a**, **Fig. S10a-c**). We used RT-qPCR to detect mRNA levels of EJC-dependent (SRSF2, 3, and 6 transcript isoforms containing PTCs) and 3’UTR EJC-independent (SMG1, 5, 6, 7, and UPF2 transcripts) NMD targets. We verified that UPF1 knockdown and rescue with WT CLIP-UPF1 resulted in marked upregulation of NMD-sensitive mRNAs and subsequent restoration back to control levels, respectively (**Fig. 6b-c**, compare siNT CLIP only to siUPF1 CLIP only to siUPF1 WT).

**Figure 6.**
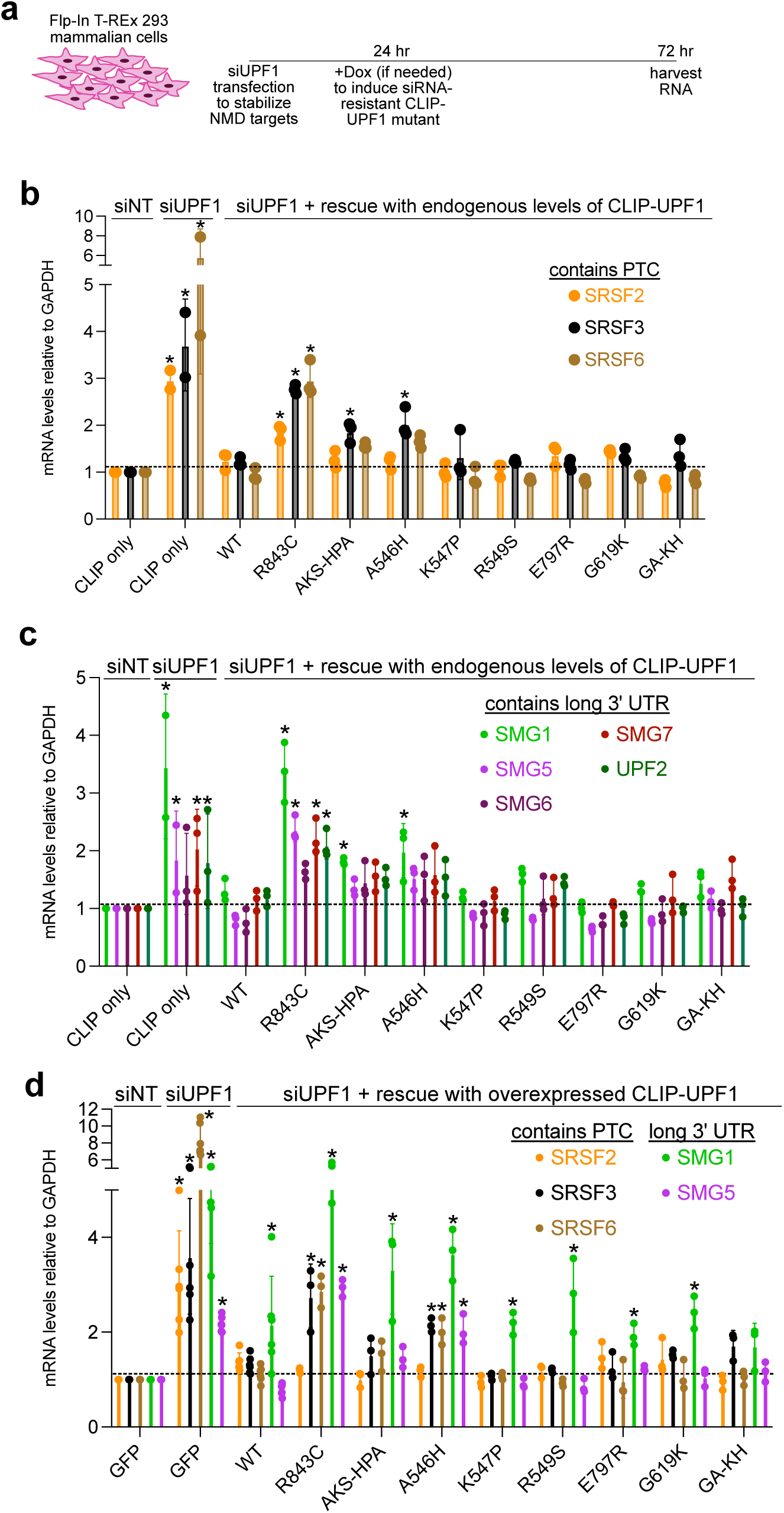
RT-qPCR reveals that UPF1 mutants with impaired helicase activity support NMD in human cells. (a) Workflow of knockdown/rescue experiments using Flp-In T-Rex 293 stably expressing CLIP-UPF1 mutants. Dox was added to induce expression of CLIP-UPF1 mutants in (d). (b-c) mRNA levels relative to GAPDH using RT-qPCR normalized to the siNT CLIP condition of EJC-dependent (b) and long 3′ UTR-dependent (c) NMD targets. siNT is non-targeting siRNA, and siUPF1 is UPF1 siRNA. CLIP-UPF1 expression was similar to endogenous UPF1 expression (**Fig. S10a**). Error bars indicate standard deviation from 3 independent experiments. (d) As in (b-c) but with CLIP-UPF1 mutants expressed ∼6-fold above endogenous UPF1 levels and normalized to the siNT GFP condition, from 3 independent experiments. Statistical significance was determined by two-way ANOVA, with comparison between the indicated conditions and siNT with CLIP only or GFP rescue (*P < 0.05, **Table S5**).

We next assayed several of the characterized mutants using this approach. As a control based on previous work, R843C was deficient in restoring NMD^16,56^, although efficiencies varied by transcript (**Fig. 6b-c**). Despite the severe biochemical defects of the two grip mutants (AKS-HPA, A546H), they were partially NMD functional (**Fig. 6b-c**). Mutants that were slower (K547P, E797R, GA-KH), faster (G619K), less processive (E797R, GA-KH), or with decoupled ATPase activity (K547P, R549S, G619K, GA-KH) restored NMD to WT or near WT levels (**Fig. 6b-c**). Importantly, GA-KH exhibited similar biochemical defects to A546H with the exception of modestly higher unwinding activity and restored ATP-stimulated dissociation from structured RNA. This dataset, in particular the ability of GA-KH to support NMD, provides strong evidence that UPF1 does not need to processively remodel mRNPs to promote NMD in human cells.

Finally, we determined whether overexpressing UPF1 mutants (**Fig. S10a,b,d**) would enhance NMD efficiency. Consistent with previous reports^36,58^, we found that overexpression of WT was not sufficient to further down-regulate canonical NMD target mRNAs (**Fig. S10e-f**, compare siNT CLIP only or GFP to siNT WT). Furthermore, mutants with impaired NMD activity under near-endogenous expression conditions (R843C, AKS-HPA, A546H) did not lead to enhanced rescue. However, these mutants led to significant upregulation of endogenous UPF1 mRNA levels in the siNT conditions (**Fig. S10d**), suggesting that an excess of faulty UPF1 causes feedback upregulation of endogenous UPF1^36,58^. Together, this quantitative characterization (**Fig. 7a**, **Table S2**) points toward a model in which UPF1 processivity, unwinding rate, and mechanochemical coupling can be significantly impaired without disrupting regulation of well-characterized NMD targets (**Fig. 7b**). These findings are consistent with models in which UPF1 uses ATP hydrolysis to sample potential substrates and perform local mRNP remodeling events to promote NMD.

**Figure 7.**
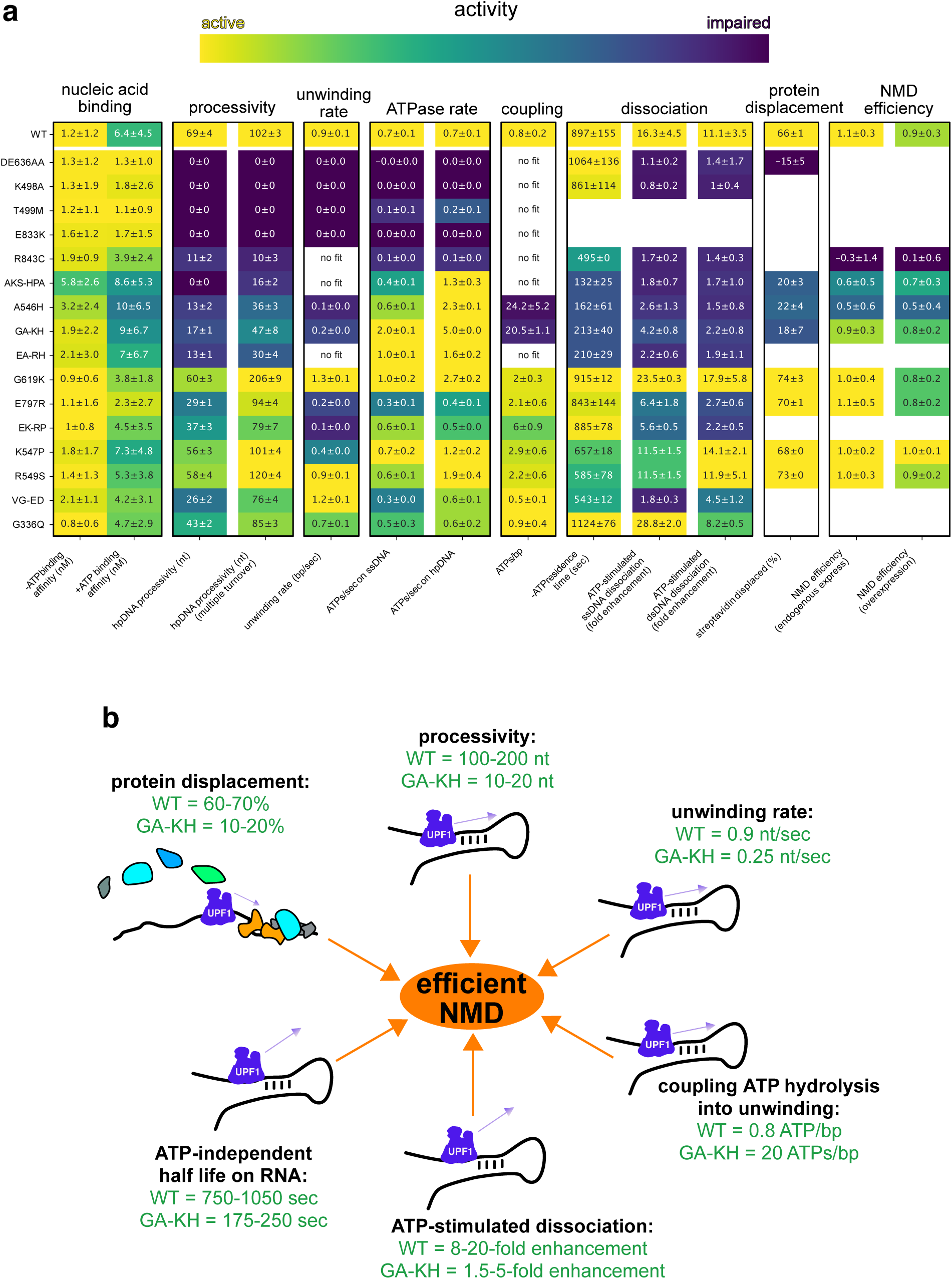
In vitro and cellular characterization reveal UPF1 mutants with slow unwinding rate, poor processivity, and poor ATPase coupling, support NMD in human cells. (a) Heatmap of mutant properties quantified from >2 independent binding assays (nucleic acid binding affinity columns), single and multiple turnover unwinding assays (processivity columns; values from best-fit curves), 10 bp hpDNA single turnover unwinding assays (unwinding rate column), ATPase assays (ATPs/sec columns), FAD assays (residence time and ATP-stimulated dissociation columns), streptavidin displacement, and relative NMD rescue from cellular experiments (see Methods). ATPs/bp is calculated by dividing ATPs/sec on hpDNA by unwinding rate. ATP-stimulated dissociation is calculated by dividing -ATP half-life by +ATP half-life on 5’ or 3’ DNA overhang substrates. Colors are relative to each column. More purple indicates greater activity detriments. VG-ED is V689E/G692D, EA-RH is E797R/A546H, GA-KH is G619K/A546H, EK-RP is E797R/K547P. (b) UPF1 enzymatic activities sufficient for efficient mammalian NMD. ATP-dependent UPF1 activities assayed in this work are shown, with properties of WT and GA-KH UPF1-HD indicated, both of which support full NMD activity on well-characterized target mRNAs.

## Discussion

UPF1 ATPase activity has long been recognized as critical for NMD. However, the precise mechanisms by which UPF1 biochemical properties influence NMD remain uncertain. The quantitative high-throughput assays developed here enable the dissection of UPF1 properties including RNA binding, ATP hydrolysis, translocation, unwinding, protein displacement, and ATPase-stimulated dissociation from RNA (**Fig. 7**). Our results reconcile reports of high UPF1 processivity (**Fig. S3**)^28,41–43^ with longstanding observations that UPF1 undergoes ATP-stimulated dissociation from nucleic acids (**Fig. S7**)^17,18,20,21,25,27,38,42^. We find that UPF1 binds very tightly to nucleic acids in the absence of ATP (∼0.5 nM K_D_, **Fig. S6-7**) but that ATP hydrolysis strongly promotes UPF1 dissociation (**Fig. S7**). This behavior results in moderate processivity (< 200 bp) when duplex-destabilizing forces are not applied to the substrate (**Fig. 1**-**2**).

### Helicase-deficient UPF1 mutants support efficient mammalian NMD

By combining our battery of biochemical assays with cellular knockdown-rescue experiments to study a panel of known and novel UPF1 mutants, we were able to define biochemical requirements for UPF1 function in NMD (**Fig. 7**). We identify UPF1 mutants with ATP hydrolysis rates similar to or higher than WT UPF1, but because of reduced mechanochemical coupling, have impaired unwinding and protein displacement activities. In particular, AKS-HPA, A546H, and GA-KH mutants unwind 1 bp per ∼20-30 ATPs hydrolyzed, resulting in slow (∼0.1-0.25 bp/sec) and poorly processive (∼10-20 nt) unwinding, and ∼4-fold lower streptavidin displacement activity. These mutants nevertheless largely or fully rescue knockdown of endogenous UPF1 (**Fig. 6**-**7**).

### Butterfly versus bulldozer models of UPF1 in NMD

Biochemical studies of UPF1 have led to two divergent models for potential roles of UPF1 enzymatic activity in NMD. In one view, which we term the butterfly model, UPF1 undergoes frequent rounds of binding and ATPase-stimulated dissociation as it surveys the transcriptome for NMD targets^35^. If UPF1 spends enough time on an RNA to sense a translation termination event and scaffold assembly of decay complexes, decay may be initiated^14,23^. This view is biochemically supported by the slow unwinding rate of human UPF1 (∼1 bp/sec) in vitro (**Fig. S3a**)^28,41,42^ and the longstanding observation that UPF1 tends to release RNA upon ATP binding/hydrolysis (**Fig. S7**)^17,18,20,21,25,27,38,42^. In further support of this model, ATPase-deficient UPF1 mutants lose substrate selectivity, becoming stably bound at non-productive sites throughout the transcriptome^16,35^, implying the importance of ATPase-stimulated dissociation for substrate selection.

In the alternative view, which we term the bulldozer model, UPF1 uses ATP to processively remodel mRNPs for efficient NMD. Support for this model is derived from the ability of UPF1 to disrupt biotin-streptavidin interactions (**Fig. 5**)^28,29^, displace nucleic acid binding proteins^28^, and from single-molecule magnetic tweezers experiments that revealed UPF1 can processively travel over 10,000 nt before dissociating (**Fig. S3b**)^28,41–43^. A characterized “grip mutant” (AKS-HPA) with severely impaired processivity in magnetic tweezers assays could not carry out NMD on the DAL7 transcript in *S. cerevisiae*, suggesting high UPF1 processivity may be important for efficient NMD^41^. However, because nucleic acid substrates are under duplex-destabilizing forces in magnetic tweezers experiments, it is unclear if high processivity is retained under physiological conditions. Notably, force-dependent processivity enhancement^59^ has been reported for the UPF1-like SARS-Cov-2 Nsp13 RNA helicase^47^ along with other SF1^44–46^, SF2^49^, and SF4^48^ helicases.

As in previous reports that used magnetic tweezers^28,41–43^, we observed very high UPF1 processivity in this context (**Fig. S3b**). However, UPF1 exhibited processivity of only 10s-100s of nt in ensemble and SPRNT assays performed without duplex-destabilizing forces applied to the nucleic acid substrate (**Fig. 1**-**2**, **Fig. S1-4**). Correspondingly, ATPase-stimulated dissociation kinetics of UPF1 from nucleic acids using FAD, SUD, and PIFE assays were consistent with moderate processivity (**Fig. 3**,**5,S5**). UPF1 appears to be a striking example of force-dependent processivity enhancement, as measured processivity differs by 2-3 orders of magnitude in the presence and absence of duplex-destabilizing forces (**Fig. 1**-**2**, **Fig. S1-4**)^28,41–43^. These forces may lower the energy barrier to forward translocation of UPF1, dramatically increasing processivity^59^. If this energy barrier corresponds to a short-lived state in the ATPase cycle, unwinding rate would remain largely unchanged, as we observed (**Fig. S3a**).

### Mechanochemically decoupled UPF1 mutants support the butterfly model

Overall, our data support the butterfly model, which requires a balance between stable UPF1 engagement with target RNAs and efficient release from non-target RNAs^35^. UPF1 can achieve this balance through ATPase-mediated control of RNA binding, as UPF1 is capable of binding RNA with sub-nanomolar affinity (**Fig. S6-7**) but can readily release RNA upon ATP hydrolysis. Without ATPase activity driving dissociation from RNA, UPF1 cannot perform proper substrate selection, instead becoming non-productively locked on mRNAs across the transcriptome^16,35^. Importantly, the robust NMD activity of mechanochemically decoupled UPF1 mutants differs markedly from that of NMD-deficient mutants with low (R843C) or no ATPase activity (previously studied ATPase-dead DE636AA, K498A, and others)^14,15,17,20,21,22,23^.

We hypothesize that a key difference between ATPase-deficient UPF1 mutants and mechanochemically decoupled mutants is the ability to undergo ATPase-stimulated dissociation. AKS-HPA, A546H, and GA-KH mutants underwent ATPase-stimulated dissociation from unstructured regions (**Fig. S8-9**). However, differential behavior on structured regions may explain the greater NMD activity of GA-KH, as A546H inefficiently dissociated from structured regions, whereas ATPase-stimulated dissociation from structured regions was partially restored in the GA-KH double mutant (**Fig. 5**). While we cannot rule out the contributions of other slight biochemical differences between A546H and GA-KH (**Fig. S8**), the increased NMD activity of GA-KH is consistent with a role for ATPase-stimulated dissociation in maintaining proper NMD.

### UPF1 ATPase-mediated release from RNA as a control point for NMD

Increasing evidence suggests that mechanisms for manipulation of ATPase-stimulated dissociation of UPF1 are important for cellular control of NMD specificity. For example, the UPF1 regulatory loop that protrudes into the RNA-binding channel mediates sensitivity to protective RNA binding proteins such as PTBP1^29,36,42^. This may be due PTBP1 exploiting certain conformational states of UPF1 to promote dissociation. Conversely, activators of UPF1-mediated mRNA decay such as Staufen and histone stem loop binding protein may promote stable association of UPF1 with mRNPs by suppressing ATPase-stimulated dissociation from target mRNAs^3^. The experimental framework developed here will enable future investigation to elucidate the biochemical mechanisms of trans-acting factors on UPF1.

---

**Figure S1.**
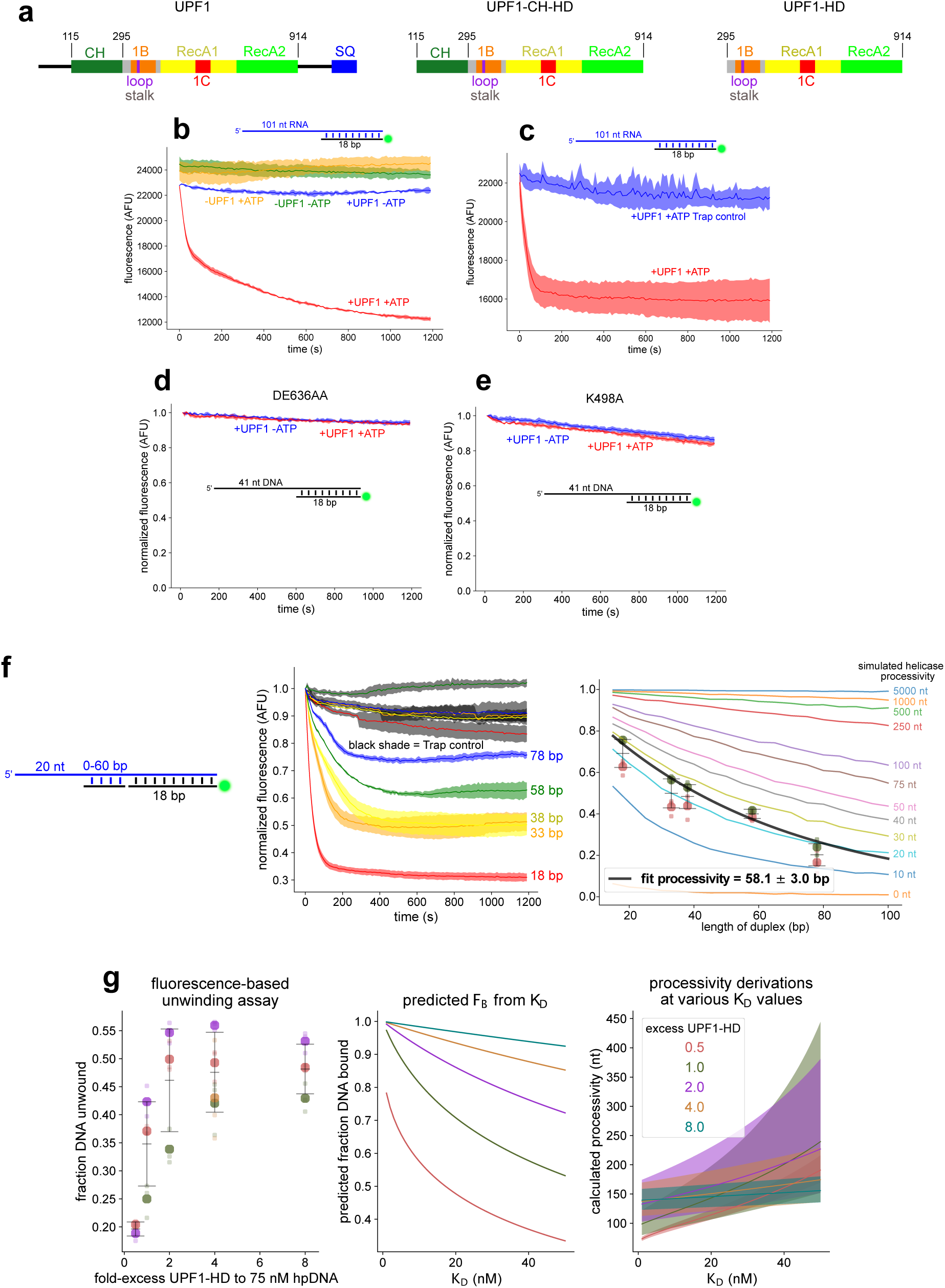
Control unwinding experiments are consistent with low to moderate UPF1-HD processivity. Related to Figure 1. (a) Schematic of UPF1 constructs. UPF1-HD was used for in vitro experiments and N-terminal CLIP-tagged UPF1 was used for cellular experiments. (b-c) Representative curves of multiple (b) or single (c) turnover unwinding of a 104 nt overhang RNA substrate from 2 independent experiments. (d-e) Representative curves of multiple turnover unwinding of a 41 nt overhang DNA substrate with ATPase-deficient UPF1-HD mutants DE636AA (d) or K498A (e). (f) Representative curves of single turnover unwinding of RNA:DNA hybrid substrates (left panel). The top strand is RNA, which was hybridized to the indicated DNA strands. Fraction unwound from 2 independent experiments (right). Fit and simulations as in Figure 1c-e. (g) Alternative processivity calculation based on a UPF1-HD titration of hpDNA unwinding. Fraction unwound of a 40 bp hpDNA substrate at different UPF1-HD concentrations (left). Each colored dot represents a separate experiment from 3 independent experiments. Small dots represent technical replicates. Predicted initial fraction of DNA bound by UPF1-HD as a function of binding affinity (K_D_) for different UPF1-HD concentrations, expressed as fold-excess to 75 nM DNA (middle). Calculation of processivity for each concentration as a function of assumed binding affinity (right, see Methods).

**Figure S2.**
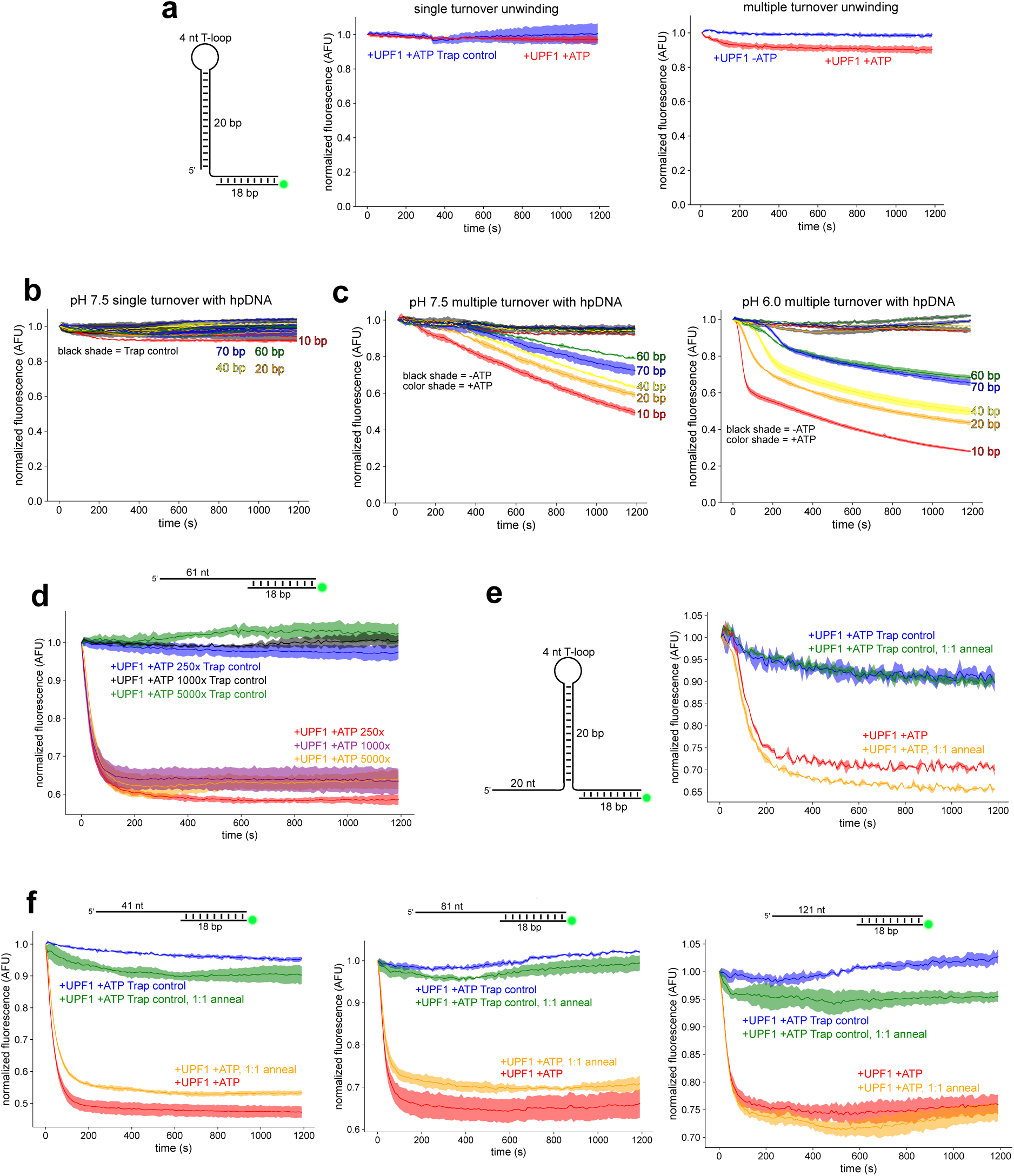
Additional control unwinding experiments reveal decreased unwinding activity at pH 7.5. Related to Figure 1. (a) Representative curves from single (left) and multiple (right) turnover unwinding measurements of a 20 bp hpDNA substrate lacking the 20 nt landing pad from 2 independent experiments. (b-c) Representative curves from single (b) and multiple (c) turnover unwinding at pH 7.5 (b; c, left) or pH 6.0 (c, right) from 2 independent experiments. (d) Single turnover unwinding at different concentrations of trap strand. (e-f) Single turnover unwinding of indicated substrates annealed using the standard protocol (see Methods) or using a 1:1 molar ratio.

**Figure S3.**
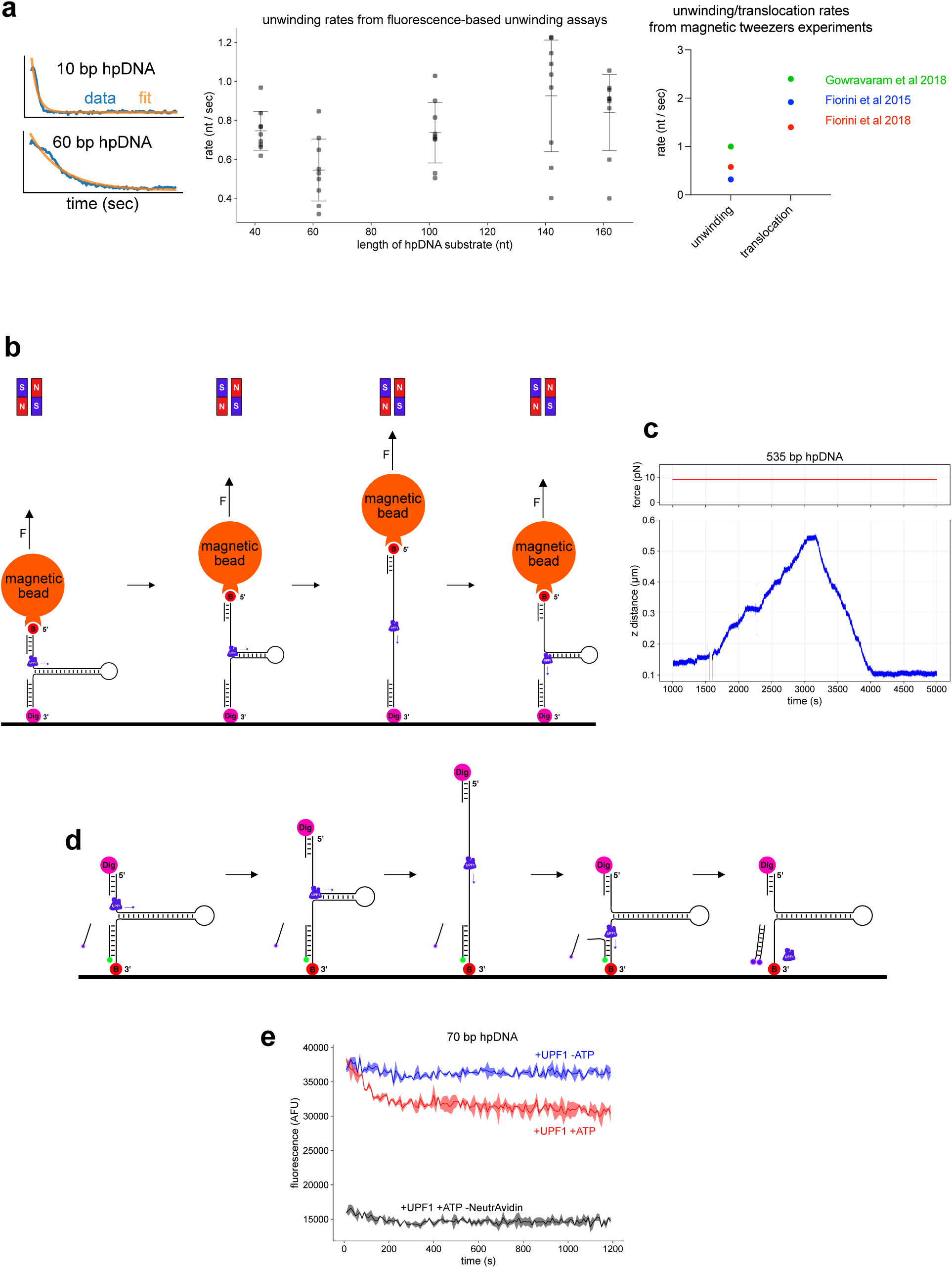
UPF1-HD processivity is higher under duplex-destabilizing force. Related to Figure 1. (a) Example fits from single turnover hpDNA unwinding curves (left) to extract unwinding rates (middle, see Methods) and comparison to rates from previous magnetic tweezers experiments (right). (b) Schematic of magnetic tweezers system. (c) Representative trace from 8 independent experiments of UPF1-HD fully unwinding and translocating through a 535 bp hpDNA substrate under 7-9 pN force. (d) Schematic of fluorescence-based unwinding assay on substrates tethered to a glass well. (e) Representative trace from 4 independent tethered unwinding experiments. The -NeutrAvidin control indicates the signal was not due to non-specific substrate binding to the surface.

**Figure S4.**
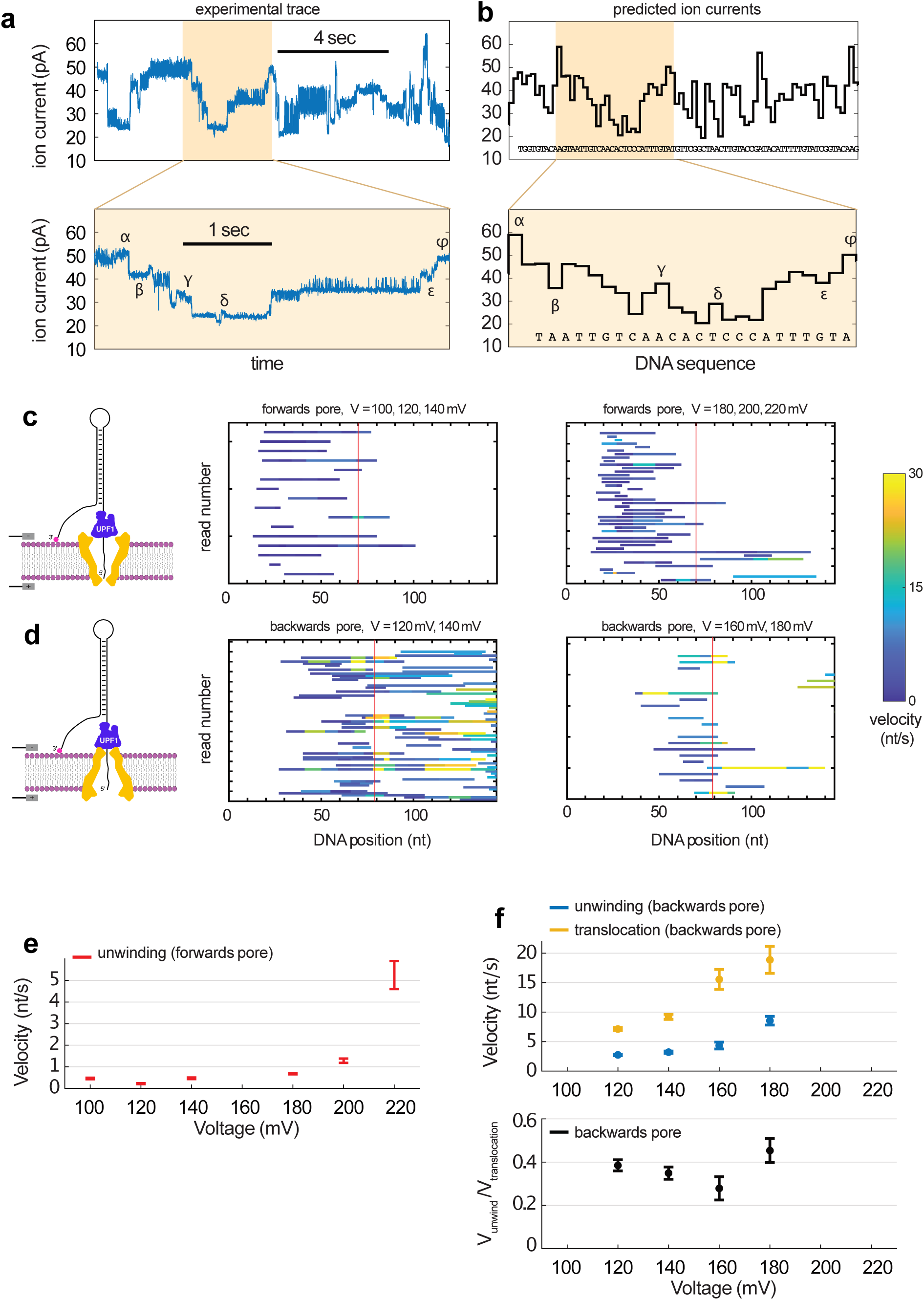
UPF1-HD velocity is force-dependent in SPRNT unwinding assays. Related to Figure 2. (a) Example UPF1-HD unwinding event (top) and zoom-in of the tan shaded area (bottom). (b) Predicted currents corresponding to experimental trace in (a). Greek letters indicate high confidence alignment of current between experimental and predicted current. (c-d) Alignment of UPF1-HD unwinding/translocation events at the indicated voltages in a forwards (c) or backwards (d) pore setup. Each read is a single event. Vertical red line indicates transition from dsDNA unwinding to ssDNA translocation. (e) Unwinding velocities in the forwards pore setup at the indicated voltages. Translocation velocities were not plotted because UPF1-HD rarely made it through the T-loop before dissociating. (f) Unwinding and translocation velocities at the indicated voltages (top) and relative unwinding/translocation velocities (bottom) in the backwards pore setup.

**Figure S5.**
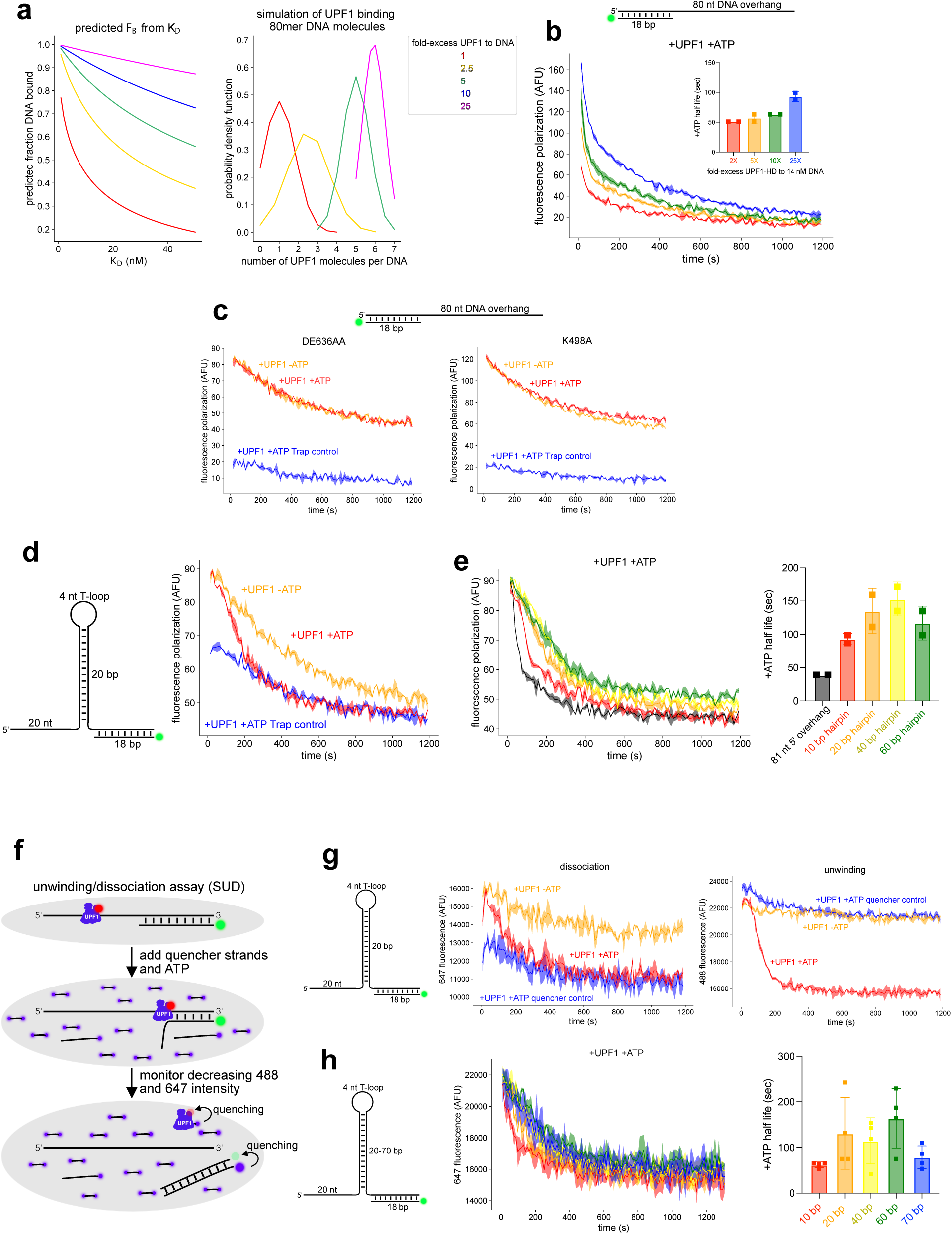
Other assays measuring in vitro dissociation of UPF1-HD are consistent with low UPF1-HD processivity. Related to Figure 3. (a) Predicted initial fraction bound by UPF1-HD as a function of binding affinity (K_D_) for different ratios of UPF1 to DNA (left). Simulations calculating the probability distribution of the number of UPF1 molecules bound to an 80 nt strand at different concentrations of UPF1-HD, assuming a 10 nt binding footprint (right). Note, the 10 and 25 conditions are perfectly superimposed. (b) Representative curves from FAD of UPF1-HD at indicated concentrations. For clarity, only +ATP curves are shown. Inset shows half lives. (c) Representative curves from FAD with a 3′ overhang substrate and DE636AA or K498A. (d) FAD assay with a 20 bp hpDNA substrate. (e) FAD assay with substrates of indicated lengths. For clarity, only +ATP curves are shown. Quantification of half lives (right). (f) Schematic of simultaneous unwinding and dissociation (SUD) assay. (g) Representative curves of SUD measuring UPF1-HD-647 dissociation from (left) and unwinding of (right) a 20 bp hpDNA substrate from 3 independent experiments. Quencher control reactions were performed by adding 250-fold excess quencher strand prior to UPF1-HD addition. (h) Representative dissociation curves of SUD with 5 hpDNA substrates from 2 independent experiments. For clarity, only +ATP curves are shown. Quantification of half lives (right).

**Figure S6.**
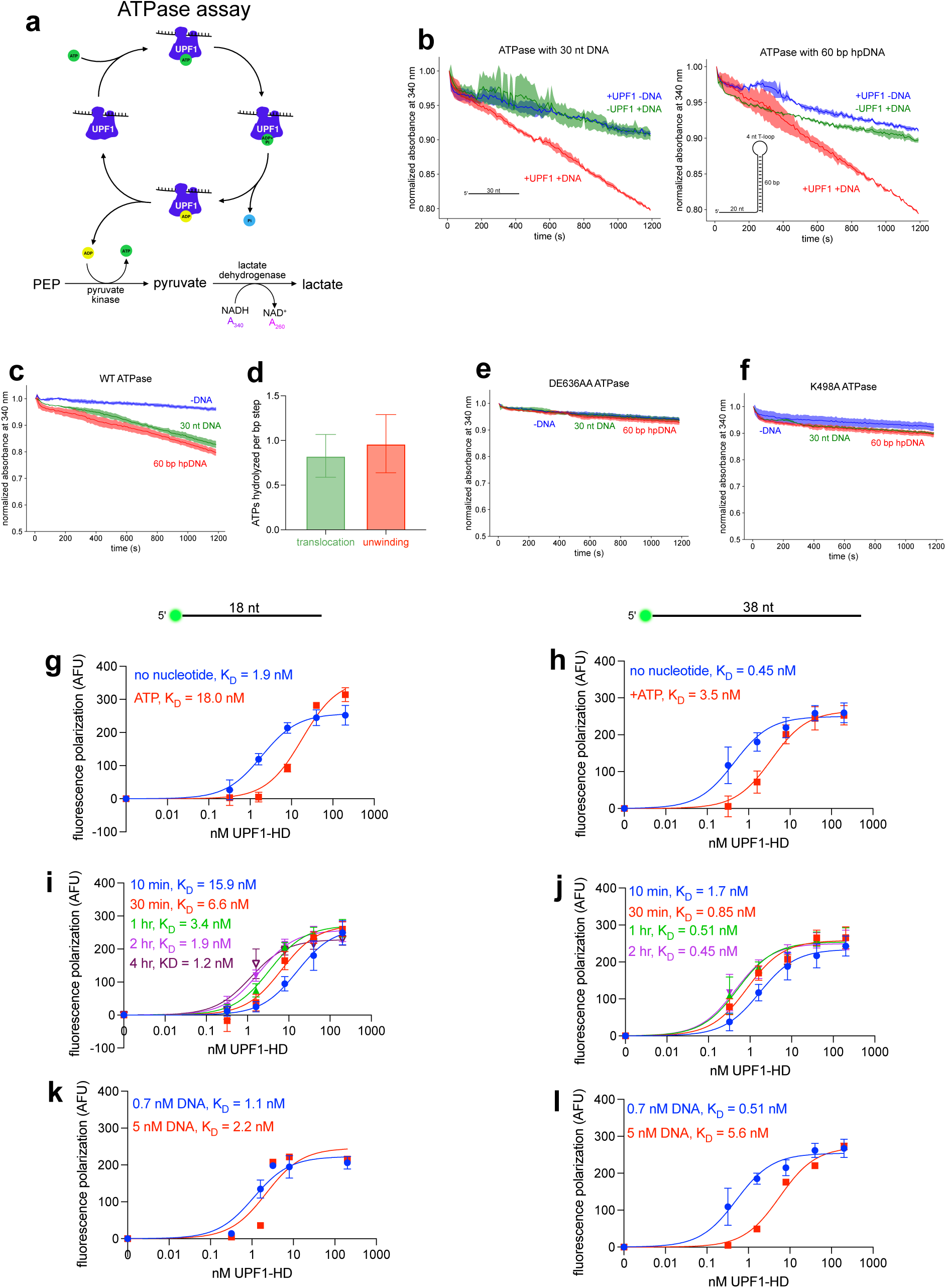
UPF1-HD exhibits sub-nanomolar nucleic acid binding affinity and tight coupling of ATP hydrolysis with unwinding and translocation. Related to Figure 3. (a) Schematic of NADH-coupled ATPase assay. (b-c) Representative curves of ATPase activity of UPF1-HD with indicated substrates from 3 independent experiments. (d) Comparison of ATPase rates to translocation and unwinding rates. (e-f) Representative ATPase assays with DE636AA (e) or K498A (f). (g-h) Binding affinities from fluorescence anisotropy binding assays with an 18 nt (g) or 38 nt (h) DNA oligonucleotide from 3 independent experiments. (i-j) Evidence that increased incubation times allow binding reactions to reach equilibrium, using 18 nt (i) or 38 nt (j) DNA oligonucleotides from 2 independent experiments. (k-l) Lowering substrate concentration reveals higher binding affinity with 18 nt (k) or 38 nt (l) DNA oligonucleotides.

**Figure S7.**
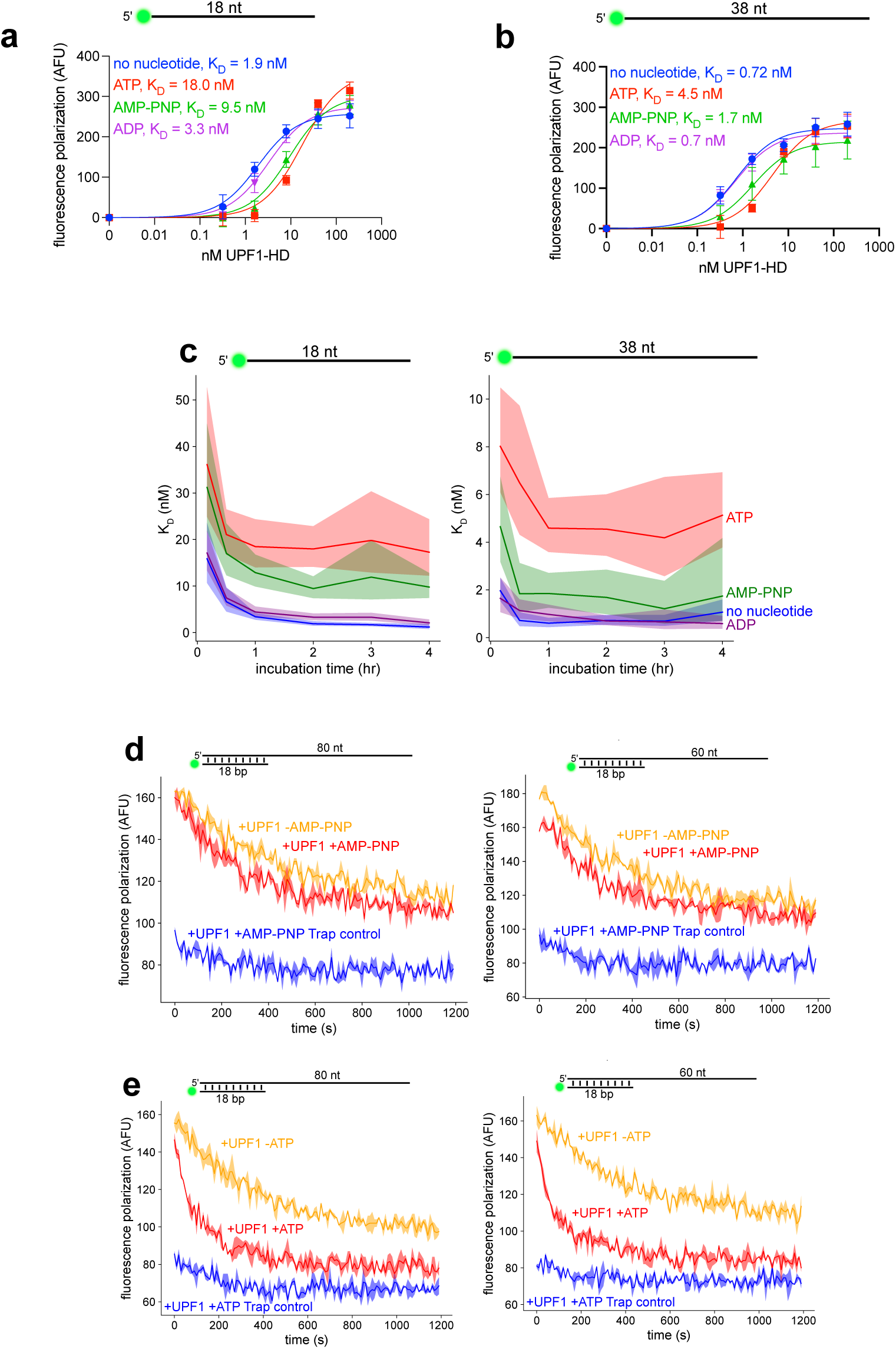
ATP binding reduces UPF1-HD binding affinity, but UPF1-HD dissociation is driven primarily by ATP hydrolysis. Related to Figure 3. (a-b) Representative binding affinities at 2 hr incubation with 1 mM of the indicated nucleotides and 18 nt (a) or 38 nt (b) DNA oligonucleotides. (c) As in (a-b), from 3 independent experiments at the indicated time points. (d-e) FAD of UPF1-HD with AMP-PNP (d) or ATP (e) with the indicated substrates, performed at 29°C from 2 independent experiments.

**Figure S8.**
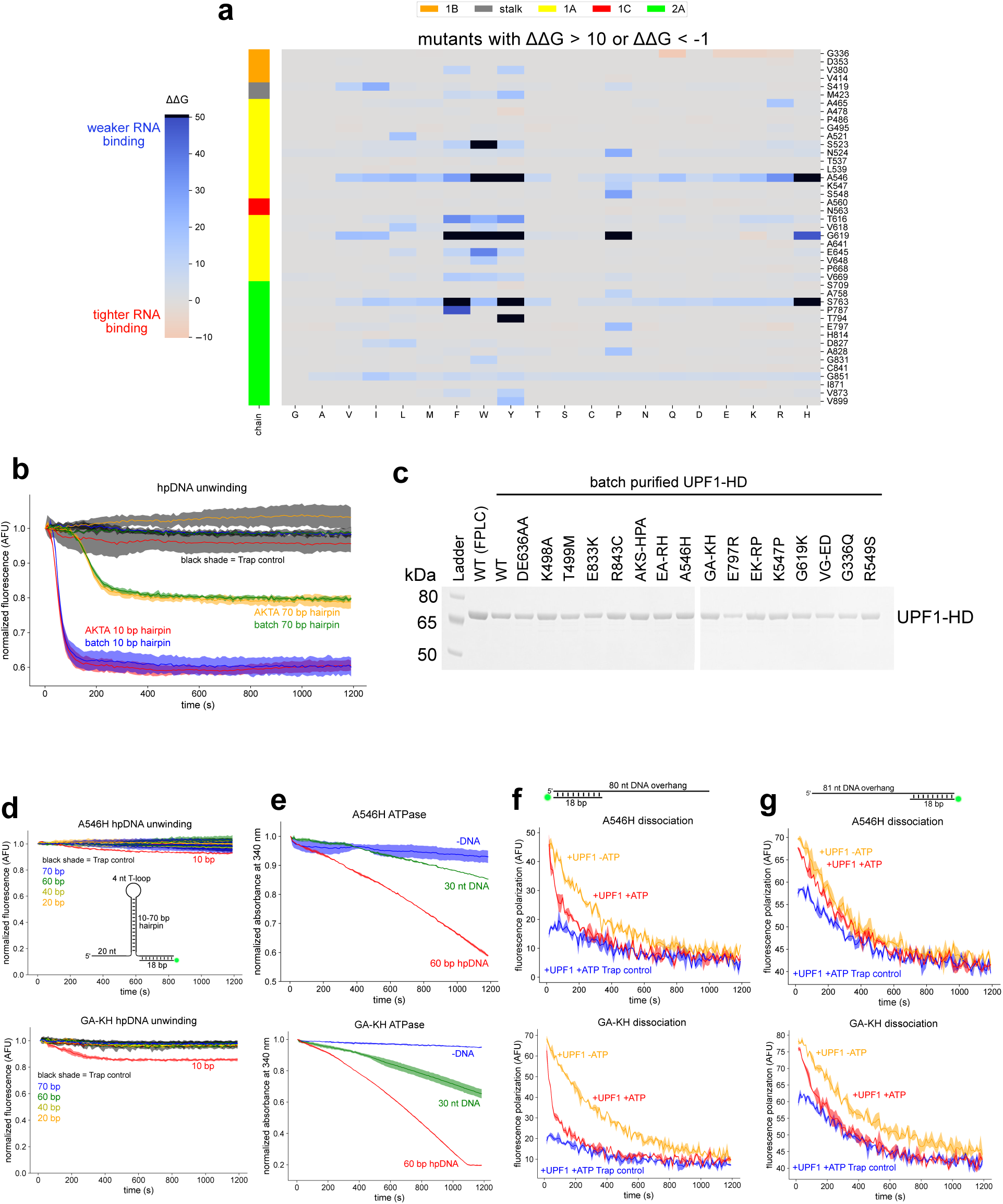
Other biochemical characterization of mutants from the Rosetta-Vienna ΔΔG Method screen. Related to Figures 4-5. (a) Results from the screen described in Figure 4a, filtering for residues exhibiting at least 1 mutation with ΔΔG > 10 or ΔΔG < -1 (units in kcal/mol). The average of 20 in silico comparisons is shown. Black boxes represent ΔΔG > 50. (b) Single turnover unwinding by UPF1-HD purified by batch purification or more stringent FPLC purification using an AKTA Protein Purification System. (c) Coomassie-stained NuPAGE 4-12% Bis-Tris Mini Protein Gel of AKTA purified WT UPF1-HD and batch purified UPF1-HD proteins. (d-g) Single turnover unwinding of hpDNA (d), ATPase (e), FAD with 3’ overhang DNA (f), and FAD with 5’ overhang DNA (g) assays using A546H (top panels) or GA-KH (bottom panels), from 2 independent experiments.

**Figure S9.**
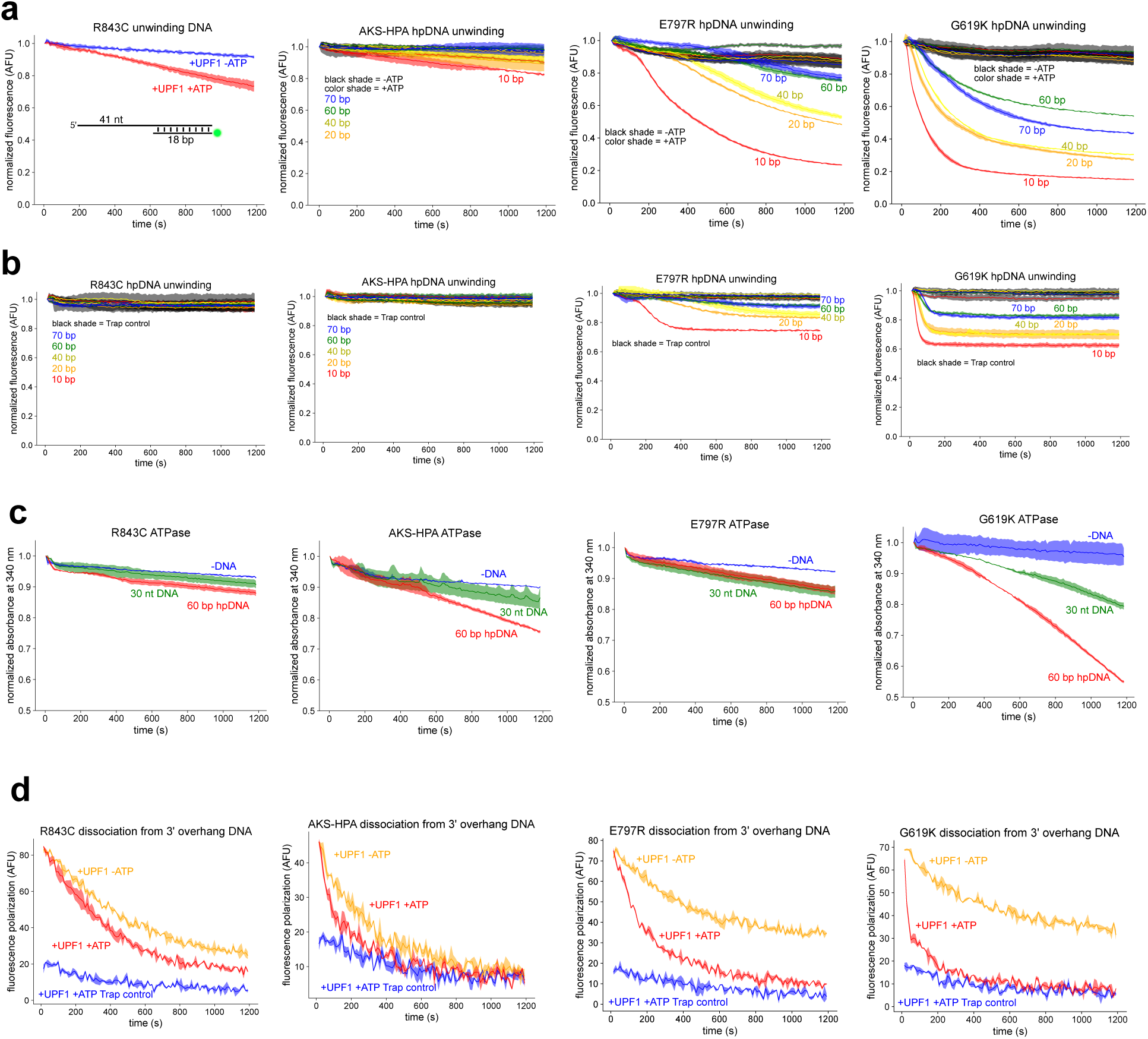
Unwinding, ATPase, and dissociation of other UPF1-HD mutants with pronounced biochemical abnormalities. Related to Figure 5. (a) Multiple turnover unwinding with the indicated substrates using R843C, AKS-HPA, E797R, and G619K UPF1-HD mutants. (b) As in (a) but under single turnover conditions. (c) ATPase assays with the indicated substrates using R843C, AKS-HPA, E797R, and G619K UPF1-HD mutants. (d) FAD assay with an 80 nt 3’ overhang DNA substrate using R843C, AKS-HPA, E797R, and G619K UPF1-HD mutants.

**Figure S10.**
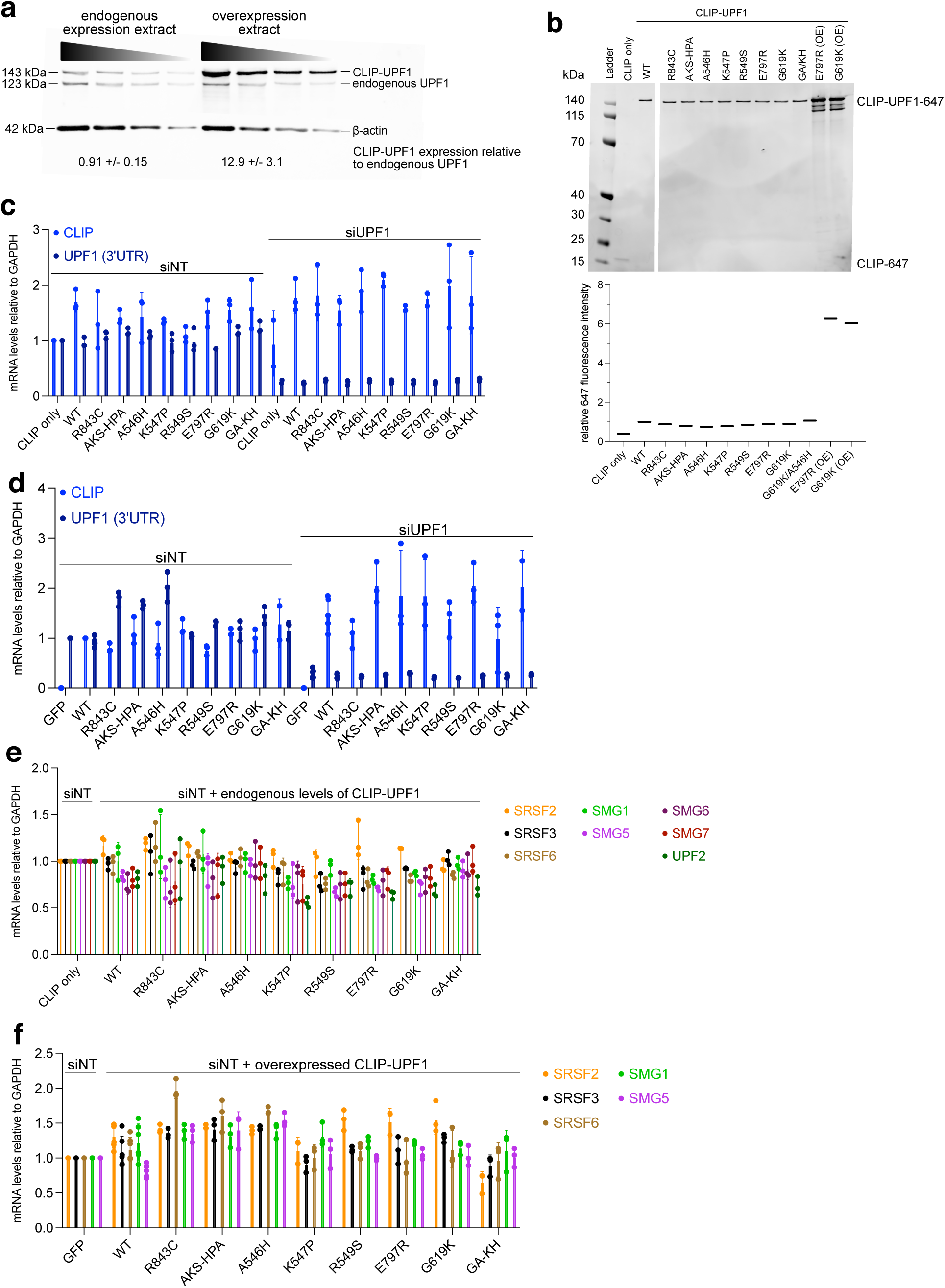
CLIP-UPF1 mutant cell line characterization reveals overexpression of mutants with ATPase-mediated dissociation defects upregulates endogenous UPF1 levels. Related to Figure 6. (a) Western blot showing levels of ectopically expressed CLIP-UPF1 compared to endogenous UPF1 levels. β-actin serves as a loading control. (b) NuPAGE 4-12% Bis-Tris Mini Protein Gel of protein extracts from CLIP-UPF1 mutant cell lines labeled with benzylcytosine-647 (top). Last 2 lanes are for comparison to overexpression cell lines. Quantification of bands (bottom). (c) CLIP tag and UPF1 3′ UTR mRNA levels relative to GAPDH using RT-qPCR normalized to the siNT CLIP condition. Error bars indicate standard deviation from 3 independent experiments. siNT is not-targeting siRNA, and siUPF1 is UPF1 siRNA. (d) As in (c) but for overexpression cell lines, and CLIP tag mRNA levels are normalized to the siNT WT condition. (e-f) siNT conditions related to Figure 6b-d for endogenously expressed (e) or overexpressed (f) CLIP-tagged UPF1 mutant cell lines.

## Methods

### Oligonucleotides and chemicals

All oligonucleotides were purchased from IDT and resuspended in TE. Biotin-BSA (Thermo Scientific 29130) was dissolved to 2 mg/mL in water, aliquoted and stored at -20°C. NeutrAvidin (Thermo Scientific 31002) was dissolved to 5 mg/mL in water, aliquoted and stored at -20°C. Alexa Fluor 647-labeled Streptavidin (Invitrogen) was dissolved to 2 mg/mL in PBS, aliquoted and stored at -20°C protected from light. Anti-Digoxigenin (Roche 11333089001) was dissolved to 0.2 mg/mL in PBS, aliquoted and stored at -20°C. Doxycycline (Sigma D9891) was dissolved to 1 mg/mL in water and stored at 4°C protected from light.

### UPF1-HD protein expression and purification

Protein expression and FPLC purification of UPF1-HD protein with N-terminal 6xHis and calmodulin binding peptide tag was performed as previously described^29^. For batch purification, after 2-3 passes through a French Press at 1000 psi, extracts were filtered through a 0.44 µm filter, rotated at 4°C for 1.5-2 hr in 0.75 mL equilibrated Ni-NTA Agarose (Qiagen), followed by packing pre-equilibrated Poly-Prep Chromatography Columns (Bio-Rad). Columns were washed with 10 mL binding buffer (1.5X PBS pH 7.5, 225 mM NaCl, 1 mM MgOAc, 0.1% NP-40, 10% glycerol) supplemented with 20 mM imidazole, followed by an additional 2.5 mL binding buffer supplemented with 50 mM imidazole, and proteins were eluted in 500-600 µL binding buffer supplemented with 500 mM imidazole. Dialysis and flash freezing was done as previously described^29^. UPF1-HD was diluted in protein dilution buffer (100 mM NaPO_4_ pH 7.2, 150 mM NaCl) for day-of use.

### In vitro transcription and substrate annealing for unwinding, dissociation, and PIFE assays

To generate RNA substrates, PCR using custom primers (**Table S3**), in vitro transcription, and quantification were performed as previously described^29^, except using T7 RNA Polymerase and pyrophosphatase generously provided by the Ferré-D’Amaré lab. Annealing of oligonucleotide substrates (**Table S3)** was also performed as previously described in 500 µL total volume^29^. Where indicated, a 1:1 substrate strand to fluorescent strand ratio was used in lieu of the standard 11:7 ratio. Annealed substrates were electrophoresed on native Novex TBE Gels (ThermoFisher Scientific) to confirm the absence of unannealed fluorophore strands.

### Fluorescence-based unwinding assays

Multiple turnover and single turnover unwinding assays were performed as previously described^29^. Briefly, 82.5 nM UPF1-HD was pre-bound to 75 nM fluorescent substrate, then 75 µM unlabeled trap strand and 562.5 nM quencher strand were added immediately before loading into the plate reader. 16 µL of 5 mM ATP (2 mM final) was injected into each well and fluorescence of the 40 µL reaction was immediately measured over time. For pH 7.5 unwinding, 20 mM Tris-HCl pH 7.5, 75 mM KOAc, 3 mM MgCl_2_, and 1 mM freshly prepared DTT were used. Measurements were performed using Tecan Spark or CLARIOstar Plus plate readers (BMG Labtech). As a single turnover control, 75 µM unlabeled trap strand was added before UPF1-HD addition. Unlabeled trap strand was omitted in multiple turnover unwinding reactions.

### Processivity calculations and unwinding data fitting

There is a competition between taking a step forward (*p_for_*) and a step off the nucleic acid (*p_off_* = 1 − *p_for_*). To calculate processivity on hpDNA, the probability of unwinding, p(u) is:

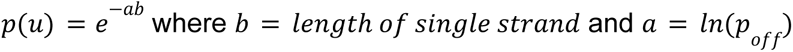

To calculate processivity on single-stranded substrates, the 18 bp duplex region was omitted for simplicity. The probability of getting to the duplex region, p(on) is:

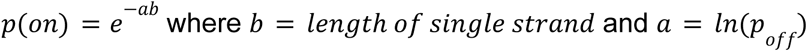

To account for initial random binding along the single-stranded region:

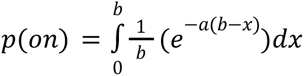

After solving the integral:

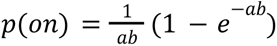

After fitting the data to the above equations with a custom python script, 𝑎 was extracted, and processivity was solved for using *processivity* = 1 − *a*. Processivity is a number between 0 and 1, but in order to calculate the average number of steps taken before dissociation^40^ (referred to in the text as processivity for simplicity):

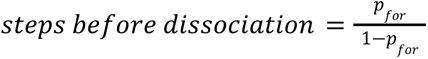

### Helicase processivity simulations

A custom python script was used to simulate helicases with differing processivities. For example, a helicase that takes 200 steps before dissociating will have a processivity of 0.995 according to the above equation. Simulating this helicase on a substrate of length 100 entails randomly selecting a number 100 times between 0 and 1. If every number is less than 0.995, that event counts as unwound. Performing this operation 1000 times will give an estimate of the fraction unwound. Iteration through different substrate lengths and processivities generated a graph of the fraction unwound against different substrate lengths (**Fig. 1c-e**, **Fig. S1f**).

### Unwinding rate data fitting

Single turnover unwinding assay results were generally fit to a single or double exponential function. A custom python script was used to fit single turnover unwinding curves normalized to single turnover control curves to the following equation:

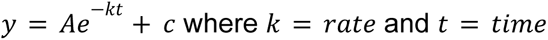

### Alternative processivity calculation

A custom python script was used to first calculate the fraction of UPF1-HD bound to nucleic acids at different concentrations using:

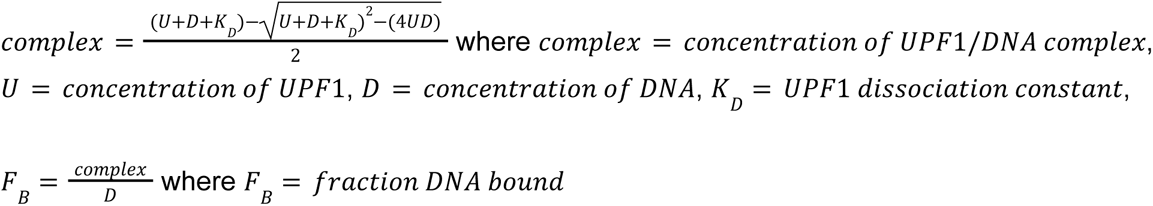

The fraction of 40 bp hpDNA (102 nt total omitting 20 nt landing pad) unwound at different UPF1-HD concentrations was then used to calculate processivity using:

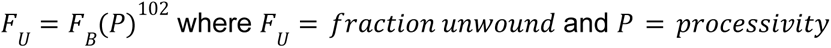

Therefore, 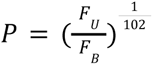

### Magnetic tweezer substrate generation

DNA hairpin substrates were created as previously described using custom oligonucleotides (**Table S3**)^61^. The surface-proximal handle was annealed to a 75 nt strand, and the bead-proximal handle was annealed to a 50 nt strand to create a 5′ 25 nt UPF1-HD loading region. The hairpin body was created via PCR of a 495 bp fragment of lambda DNA to generate a 535 bp hairpin stem.

### Magnetic tweezer force-based unwinding assay

Experiments were performed as previously described^61^. Briefly, a 30 µL reaction containing 50 pM hpDNA substrate and 6.67 ng/µL anti-digoxigenin was added to a KOH-cleaned and assembled flow cell with polystyrene beads melted on the surface^61^. After incubating at 4°C overnight, 200 µL wash buffer (PBS supplemented with 0.3% BSA and 0.02% Tween-20) was flowed through the flow cell, which was then placed on a PicoTwist magnetic tweezer apparatus and allowed to settle for 30 min. Next, 5 µL of 10 mg/mL Dynabeads MyOne Streptavidin T1 magnetic beads (Invitrogen 65601) were washed twice in 200 µL bead wash buffer (10 mM Tris-HCl pH 7.0, 2 M NaCl, 1 mM EDTA), resuspended in 200 µL wash buffer, and sonicated for 5 min, after which 40 µL was flowed into the flow cell and incubated for 5 min. Next, 1-2 mL wash buffer was flowed through until most unbound beads were gone, after which 0.5-1 mL UPF1 wash buffer (20 mM Tris-HCl pH 7.5, 75 mM KOAc, 3 mM MgCl_2_, 1 mM DTT, 1% BSA, 0.02% Tween-20) was flowed through, and tethers were located and calibrated. Only tethers that extended 0.4-0.5 µm as the force was increased from 7-9 pN to 20-22 pN were used in experiments. 200 µL UPF1 wash buffer supplemented with 20 nM UPF1-HD was then passed through 3 times, waiting 3-4 min after each addition. A final 200 µL was passed through, incubated for 10 min, then 100 µL UPF1 wash buffer supplemented with 2 mM ATP was added to the flow cell, 25-50 µL of which was passed through to initiate UPF1-HD unwinding.

### Tether unwinding substrate generation

After annealing four oligonucleotides (hairpin strand, digoxigenin strand, biotin strand, Alexa Fluor 488 strand, **Table S3**) in an 80 µL reaction, 7.5 µL T4 DNA Ligase in T4 DNA Ligase Buffer (NEB) was added, and the reaction was incubated at room temperature for 1 hr, followed by heating at 65°C for 10 min and cooling to room temperature for 1 hr on a tube block. Reactions were split in two and passed through two CHROMA SPIN+TE-100 columns (Takara 636072), following the manufacturer’s protocol. Denaturing (Novex TBE-Urea) and native (Novex TBE) gels were run for quantification and quality checks.

### Tether unwinding assay

Biotin-BSA was dissolved to 1 mg/mL in tether buffer (50 mM Tris-HCl pH 7.5, 50 mM NaCl). 5 µL was added to the center of a well in a 24 well glass bottom plate (Cellvis P24-1.5H-N) and incubated for 15 min. After 3 washes with 5 µL tether buffer, 5 µL of 1 mg/mL NeutrAvidin (in tether buffer) was added and incubated for 15 min to bind exposed biotin moieties. After 3 tether buffer washes, the 5 µL spot was washed 3 times with passivation buffer (2% BSA, 1 mM DTT, and 2 mM MgOAc). Next, 200 nM hairpin substrate pre-incubated with 200 nM UPF1-HD at room temperature in passivation buffer for 10-15 min was added to the spot, incubated for 15 min, and washed 3 times with passivation buffer to remove unbound substrates. Lastly, 15 µL passivation buffer supplemented with 2 mM ATP and 600 nM of a 38 nt quencher strand was added and Alexa Fluor 488 fluorescence was measured over time, as in solution fluorescence-based unwinding assays at 37°C in a CLARIOstar Plus plate reader (BMG Labtech).

### Single-molecule picometer-resolution nanopore tweezers (SPRNT) substrate generation

All DNA oligonucleotides for SPRNT experiments were ordered from panoligo at Stanford. Hairpins were formed by rapid annealing, in which 1 µM DNA was heated to 90°C then cooled to 4°C over 4 min.

### SPRNT unwinding and translocation assays

SPRNT assays were performed as previously described^52^. 4.5 µM UPF1-HD was incubated with 1 µM DNA at 37°C, of which 1 µL was added to a 100 µL reaction volume. Experiments were run at 1 mM or 2.5 mM ATP supplemented with 10 mM MgCl_2_ and 10 mM DTT. MspA nanopores insert in a random orientation, and because MspA is a non-ohmic pore, the pore orientation is determined by measuring the current-voltage response of the pore. At 500 mM KCl, a forwards pore has a current of 180±10 pA at 180 mV and -220±10 pA at -180 mV. Conversely, a backwards pore has a current of 220±10 pA at 180 mV and -180±10 pA at -180 mV.

### Processivity analysis from SPRNT measurements

The ionic current response to DNA sequence for forwards and backwards pores for enzymes such as Hel308 helicase and Phi29 DNA polymerase has been previously measured^50,51^. This response allows for prediction of the ionic current pattern from the known DNA sequence. Alignments were done using an in-house GUI to identify features in the ionic current trace as in **Fig. S4a-b**, where the greek letters indicate features that are readily identified from the experimental data. The mean helicase velocity taken over a given set of traces is calculated as: 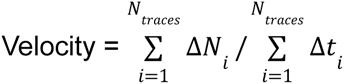 where Δ*N_i_* is the nt distance traveled by a single UPF1 before dissociation and Δ*t_i_* is the total time from the start of the read to the end.

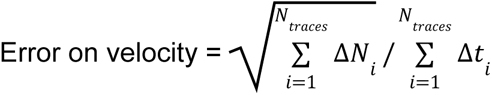

Processivity was calculated by binning the distribution of Δ*N_i_* into 10 nt intervals and fitting the histogram (omitting the first bin due to difficulty identifying short traces and the last bin due to integration of reads longer than the DNA length) to a single exponential function 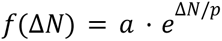 where p is the processivity in nt.

### Fluorescence anisotropy dissociation (FAD) assay

15 µL reaction buffer (10 mM MES pH 6.0, 50 mM KOAc, 0.1 mM EDTA, 2 mM MgOAc, 2 mM DTT) supplemented with 14.25 nM substrate (**Table S3**) was added to a 96 well half-area black microplate (Corning 3993), followed by addition of 3 µL of 950 nM UPF1-HD (71.25 nM final) and incubation for 10 min. Next, 3 µL of 190 µM trap strand was added (14.25 µM final), and the plate was loaded into a Tecan Spark or CLARIOstar Plus plate reader (BMG Labtech) pre-heated to 37°C. Fluorescence polarization of Alexa Fluor 488 was measured over time, beginning immediately after injection of 16 µL of 5 mM ATP (2 mM final) into wells. To obtain a baseline of unbound substrates, 14.25 µM trap strand was added prior to UPF1-HD addition, and constant low polarization over time was observed because very few fluorescent strands were protein-bound. 5-fold excess UPF1-HD was used in order to increase the dynamic range of the assay. Half-life was computed by extracting the time at which fluorescence polarization decreased halfway to the trap control baseline.

### UPF1-HD fluorescent labeling with cysteine-maleimide chemistry

Batch purified UPF1-HD was re-dialyzed in 500 mL DTT-free buffer (1.5x PBS (KD Medical), 125 mM NaCl, 0.66 mM MgOAc, 1 µM ZnSO_4_) in 0.5 mL Slide-A-Lyzer Dialysis Cassettes (Thermo Scientific 66454) at 4°C for 4 hr replacing buffer every hr. 1 mM TCEP was added to reduce cysteine residues for 10 min on ice. Next, 100 µM Alexa Fluor 647 C_2_ Maleimide (10-fold molar excess, Thermo Fisher Scientific) was added and rotated at 4°C overnight. 10 mL DTT-free dialysis buffer and 10 mL regular dialysis buffer were freshly made and cooled on ice. Zeba Spin Desalting Columns (Thermo Fisher Scientific 89882) were equilibrated following the manufacturer’s protocol. Samples were passed through 3 times in columns equilibrated with DTT-free dialysis buffer, then passed through once in columns equilibrated with regular dialysis buffer.

### Simultaneous unwinding/dissociation (SUD) assays

SUD experiments were performed in a similar manner as single turnover unwinding assays, except 250-fold excess of dual quencher (12 nt oligonucleotide dual labeled with Iowa Black RQ to quench UPF1-HD-647, **Table S3**) strand was substituted for the 1000-fold excess of unlabeled trap strand, and UPF1-HD-647 for UPF1-HD. All experiments were performed on a CLARIOstar Plus plate reader at 37°C (BMG Labtech).

### NADH-coupled ATPase assays

Assays were performed as previously described where 80 nM UPF1-HD was pre-bound to 1 µM substrate^29^ but (**Table S3**) supplemented with 1 mM DTT and using clear 96 well half area plates (Greiner Bio-one 675101). All experiments were performed on a CLARIOstar Plus plate reader at 37°C with 1 mM ATP (BMG Labtech).

### ATPase assay data fitting

Data was fit using a linear regression with the pwlf python package^62^. The slope of the line (*A*) was converted into change in NADH concentration (*c*) over time using Beer’s Law:

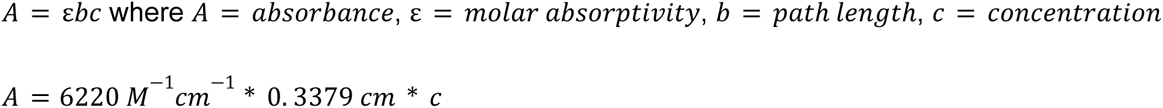

where the path length was determined by measuring the height of the reaction in the well up to the meniscus. This was divided by UPF1-HD concentration (80 nM) in solution to obtain molecules of NADH oxidized per UPF1-HD molecule per second.

### Fluorescence anisotropy binding assays

37 uL master mix (20 mM Tris-HCl pH 7.5, 75 mM KOAc, 3 mM MgCl_2_, 1 mM freshly prepared DTT, 1 mM nucleotide, and indicated concentration of Alexa Fluor 488 labeled DNA, **Table S3**) was added to a 96-well half area black plate (Corning 3993) followed by addition of 3 uL protein diluted in protein dilution buffer (100 mM NaPO_4_ pH 7.2, 150 mM NaCl) and incubated for the indicated time points. Wells were measured using the fluorescence polarization module on the Tecan Spark plate reader. Note that the apparent binding affinities observed here may be underestimated, as substrate concentrations below 0.5 nM were below the limit of detection for this assay.

### PIFE assays

18 µL reaction buffer (10 mM MES pH 6.0, 50 mM KOAc, 0.1 mM EDTA, 2 mM MgOAc, 2 mM DTT) supplemented with 75 nM DNA or 38 nM RNA substrate (**Table S3**) was added to a 96 well half-area black microplate (Corning 3993), followed by addition of 3 µL of 1.1 µM or 550 nM UPF1-HD (82.5 or 41.25 nM final) and incubation for 10 min. Next, 3 µL of 1 mM or 0.5 mM trap strand was added (75 or 37.5 µM final), and the plate was loaded into a CLARIOstar Plus plate reader (BMG Labtech) pre-heated to 37°C. Cy3 fluorescence intensity was measured over time, starting immediately after injection of 16 µL of 5 mM ATP (2 mM final) into wells. To obtain a baseline of unbound substrates, 75 or 37.5 µM trap strand was added prior to UPF1-HD addition. Half-life was computed by extracting the time at which fluorescence polarization decreased halfway to the trap control baseline.

### Fluorescence-based biotin-streptavidin displacement assays

15 µL reaction buffer (10 mM MES pH 6.0, 50 mM KOAc, 0.1 mM EDTA, 2 mM MgOAc, 2 mM DTT) supplemented with 75 nM of a 3′ biotin-labeled DNA substrate (**Table S3**) was added to a 96 well half-area black microplate (Corning 3993), followed by addition of 3 µL of 2.2 µM UPF1-HD (165 nM) and incubation for 5 min. Next, 3 µL of 200 nM Alexa Fluor 647-labeled Streptavidin tetramer (Invitrogen, 15 nM final) was incubated in the reaction for 10 min, and 3 µL of 63 µM of a 12 bp biotin-quencher was added (4.725 µM final). The biotin-quencher was generated by annealing a 12 nt 5′ biotin-labeled oligonucleotide with dual quencher (see SUD assay), following the standard protocol but using 1.23-fold excess of dual quencher. After a 10 min incubation to allow the reaction to come to equilibrium, the plate was loaded into a CLARIOstar Plus plate reader (BMG Labtech) pre-heated to 37°C. Alexa Fluor 647 fluorescence intensity was measured over time, starting immediately after injection of 16 µL of 5 mM ATP into wells (2 mM final). To obtain a baseline of quenched streptavidin, 4.725 µM biotin-quencher was added prior to Alexa Fluor 647-labeled Streptavidin addition.

### In silico UPF1-HD structure preparation for mutational screen

The UPF1-HD crystal structure (PDB 2XZO^38^) was modified before performing the in silico screen. First, the terminal nucleotide was patched using rosetta/2019.42/main/database/chemical/residue_type_sets/fa_standard/patches/nucleic/rna/3p rime_phosphate.txt and subsequently minimized. The structure was further curated with BioLuminate (https://www.schrodinger.com/products/bioluminate) using the Protein Preparation Wizard tool. Missing side chains were filled in using Prime, and missing loops were filled in by referencing a curated database of known loops in the PDB. The curated structure was relaxed 100 times using rosetta/2019.42/main/source/src/apps/public/rnp_ddg/get_lowest_scoring_relaxed_models.py --relax_dir relax_reprocessed > lowest_scoring_relaxed_structures.txt, where the 20 lowest ΔΔG structures were used for downstream analysis. This structure is in a post-hydrolysis state, which maximizes the likelihood of finding important interactions with RNA at an intermediate state in the ATPase/helicase cycle.

### In silico Rosetta-Vienna RNP ΔΔG method^54^ for mutational screen

Sequences of every possible point mutation were generated using a custom Python script and subsequently split into 100 text files to parallelize computing. Custom bash scripts were run on the NIH high-performance computing system, Biowulf, to complete the Rosetta-Vienna RNP ΔΔG method^54,55^. Each mutant was run 20 times to calculate the average and standard deviation ΔΔG values.

### Site-directed mutagenesis of UPF1-HD plasmids for bacterial expression

The pET28 vector harboring UPF1-HD with an N-terminal 6xHis and calmodulin binding peptide tag was used as the template. Mutations were generated by following the Q5 site-directed mutagenesis kit protocol (NEB E0554) with Q5 Hot-Start High-Fidelity Master Mix, 10 ng template plasmid, and 500 nM of appropriate primers (designed via the NEBaseChanger website, **Table S3**). PCR was initiated at 98°C for 30 sec, followed by 25 cycles of 98°C 10 sec, T_a_ 20 sec, 72°C 3.5 min, and a final 72°C extension for 2 min. An aliquot of the 25 µL reaction was run on a 1% ethidium bromide agarose gel (or 1% E-Gel agarose gel, Thermo Fisher Scientific) to verify 1 unique PCR product band. The PCR reaction was purified using the Qiagen QIAquick PCR Purification Kit eluting in 30 µL buffer EB. 1 µL was used in a 10 µL Kinase/Ligase/DpnI (KLD, NEB) reaction and incubated at room temperature for 10-20 min. Following KLD treatment, 5 µL was transformed into either NEB 5-alpha competent *E. coli* cells or homemade DH5-alpha competent cells and plated onto LB agar plates with 30 µg/mL kanamycin (Teknova L1024). After a 37°C overnight incubation, colonies were picked into 3 mL LB supplemented with 50 µg/mL kanamycin and, after a 37°C 230 rpm incubation overnight, cultures were subjected to miniprep using Qiagen QIAprep Spin Miniprep Kit eluting in 30 µL buffer EB. Clones were sequence verified with custom primers (**Table S3**, Psomagen).

### Site-directed mutagenesis of CLIP-UPF1 plasmids for mammalian overexpression

The pcDNA5-FRT-TO vector harboring N-terminal CLIP-tagged full length UPF1 was used as the template ^63^. Mutations were generated as above, but due to the high GC content and vector size, several modifications were necessary. Phusion high-fidelity DNA polymerase in Phusion GC buffer and 3% DMSO were used to set up PCR reactions, which were initiated at 98°C for 5 min followed by 25 cycles of 98°C 1 min, T_a_ (minus 5-6°C 30 sec, 72°C 5 min, and a final 72°C extension for 10 min. If multiple bands appeared on the gel, the entire PCR reaction was rerun on the gel, bands were cut out, and purified using the Qiagen MinElute Gel Extraction Kit eluting in 10 µL buffer EB. Transformed competent cells were plated onto LB agar plates with 100 µg/mL carbenicillin (Teknova L1010), and colonies were picked into LB supplemented with 50 µg/mL carbenicillin. For transfection-grade plasmids, the NucleoSpin Miniprep kit was used (Macherey-Nagel 740490), and plasmids were eluted in 50-60 µL elution buffer. Clones were sequence verified with custom primers (**Table S3**, Psomagen).

### Cloning of CMV (−40)+1 CLIP-UPF1 plasmids for mammalian expression

The pcDNA5-FRT-TO-CLIP-UPF1 plasmids are overexpressed in mammalian cells, so a -40 kcal/mol hairpin sequence was placed 1 nt downstream of the predicted transcription start site to lower CLIP-UPF1 protein expression. The tetracycline-regulated promoter of the pcDNA5-FRT-TO sequence was replaced with the CMV promoter and hairpin sequence by cut-and-paste cloning. 1 µg of each pFRT-TO-CLIP-UPF1 mutant vector was digested with 1 µL SpeI (NEB) and 1 µL HindIII-HF (NEB) in CutSmart buffer in 20 µL at 37°C overnight. The same reaction was performed to cut out the CMV (−40)+1 sequence but with 3 µg plasmid. Restriction digests were run on 1% E-Gel EX agarose gels (Thermo Fisher Scientific), and bands were cut out. Vector digest bands were purified using the Qiagen QIAquick Gel Extraction Kit eluting in 30 µL buffer EB, whereas the CMV (−40)+1 insert sequence band was purified using the Qiagen MinElute Gel Extraction Kit eluting in 10 µL buffer EB. A 10 µL reaction containing 3 µL vector, 3 µL insert, and 1 µL T4 DNA Ligase (NEB) in T4 DNA Ligase buffer was incubated at room temperature for 30-60 min, and subsequently transformed into DH5-alpha cells and cultures were grown, purified, and sequence verified as above.

### Mammalian stable cell line generation

T-Rex-293 cells (Invitrogen) were cultured and maintained as previously described^63^ in DMEM (Gibco 11-965-092) supplemented with 10% FBS (Gibco) and 1x penicillin/streptomycin/L-glutamine (Gibco 10378016) at 37°C and 5% CO_2_. To generate stable lines, 500,000 cells were split into each well in a 6-well plate (Corning 353046). 16-18 hr later, cells were transfected using the Lipofectamine 3000 Reagent (Thermo Fisher Scientific) with 200 ng pcDNA5-FRT-TO-CLIP-UPF1 and 2.25 µg pOG44 Flp-Recombinase Expression Vector (or pcDNA3.1 as a control). Media was replaced 6-7 hr post-transfection. Cells were transferred into 10 cm plates (Corning 353003) 2-3 days post-transfection, and media was replaced after 4-16 hr with media supplemented with 100 µg/mL Hygromycin B (Invitrogen) to start selection. Once visible colonies started forming, cells were transferred into 15 cm plates (Corning 353025). After 1-2 splits, cells were trypsinized (Gibco 12605), transferred into 15 mL tubes, centrifuged at 1000 *g* for 1 min, washed in DPBS (Gibco 14190), and resuspended in 5 mL freeze media (DMEM supplemented with 30% FBS and 10% DMSO), then aliquoted into cryogenic vials, slowly frozen at -80°C in CoolCell Alcohol-Free Cell Freezing Containers (Thomas Scientific) for at least 3 days, then transferred onto dry ice then into liquid N_2_ tanks for long-term storage.

### Protein extraction from cell lines and CLIP tag labeling

Cells in 12-well plates (Corning 353043) were lysed in 300 µL Passive Lysis Buffer (Promega) and centrifuged at maximum speed for 1-2 min at 4°C. The supernatant was transferred to pre-cooled tubes, of which 10 µL was mixed with 150 µL Pierce 660nm Protein Assay Reagent (Thermo Scientific) in a clear 96-well plate (Greiner Bio-one 655101). After a 5 min incubation on a rocker, absorbance at 562 nm was measured on a Tecan Infinite M200 Pro plate reader. Following quantification, a 25 µL reaction containing 0.2 mg/mL extract, 1 mM DTT, and 8 µM BC-647 (CLIP-Surface 647, NEB) in a base solution of Passive Lysis Buffer was rotated overnight at 4°C to label CLIP-tagged proteins. An aliquot of this reaction was mixed with 8 mM DTT and NuPAGE LDS Sample Buffer (Invitrogen), heated at 70°C for 5 min, then loaded on a NuPAGE 4-12% Bis-Tris Mini Protein Gel in MOPS buffer with 500 µL NuPAGE Antioxidant (Invitrogen) in the inner chamber. The gel was run at 100 V for 10 min then 200 V for 40 min, imaged using an Amersham Typhoon (GE), and quantified using Fiji.

### siRNA treatment and cell harvesting

6 µL of 20 µM siRNA (siUPF1: sense = CUACCAGUACCAGAACAUAtt, antisense = UAUGUUCUGGUACUGGUAGgc, siNT: AN2) were added to 6-well plates, then 1.5 mL master mix (1.5 mL Opti-MEM plus 5.25 µL Lipofectamine RNAiMAX) was added and incubated for 30-50 min. During the incubation, cells were trypsinized, counted, and diluted to 2×10^5^ cells/mL with media containing 20% FBS, of which 1.5 mL was added to each well. 24 hr later, media was replaced with media supplemented with 200 ng/mL doxycycline to induce expression where applicable. 72 hr post-siRNA transfection, cells were washed in DPBS and harvested by resuspending in 500 µL TRIzol Reagent (Invitrogen) and stored at -80°C.

### Western blotting

Following cell lysis and quantification as above, extract was equilibrated with Passive Lysis Buffer (Promega), and aliquot was mixed with 8 mM DTT and NuPAGE LDS Sample Buffer (Invitrogen), heated at 70°C for 5 min, then loaded on a 1.0 mm NuPAGE 4-12% Bis-Tris Mini Protein Gel in MOPS buffer with 500 µL of NuPAGE Antioxidant (Invitrogen) in the inner chamber. The gel was run at 100 V for 10 min then 200 V for 40 min, then transferred to a 0.45 µm nitrocellulose membrane (BioRad 1620115) and transferred using the XCell II Blot Module (Invitrogen). Following transfer, membranes were cut, then incubated in 5 mL Blocking Buffer (Rockland MB-070) for 1 hr at room temperature. Next, membranes were incubated overnight at 4°C with 5 mL primary antibodies diluted in Blocking Buffer. 1:5000 goat anti-Rent1 (Bethyl A300-038A) was used for UPF1 detection, and 1:5000 mouse anti-beta actin (Cell Signaling 3700) was used for beta actin detection. Next, membranes were washed 3 times in 5 mL TBS supplemented with 0.1% Tween-20, then incubated for 90 min at room temperature with 5 mL of 1:10,000 of either donkey anti-goat AlexaFluor 680 or goat anti-mouse AlexaFluor 680 secondary antibodies. Membranes were subsequently imaged using an Amersham Typhoon (GE) and quantified using Fiji.

### RNA extraction, cDNA synthesis, RT-qPCR

RNA was extracted from mammalian cells by following the TRIzol Reagent manufacturer protocol but using 1 µL GlycoBlue (Invitrogen AM9515) to visualize nucleic acid pellets. DNase treatment using RQ1 DNase (Promega) was performed, and RNA was re-extracted using acid-phenol and precipitated with standard ethanol precipitation. DNase treatment using 1 µL Shrimp dsDNase (Thermo Scientific EN0771) in a 10 µL reaction for 5 min at 25°C then 55°C for 5 min was also used, followed by cleanup with 18 µL RNAclean XP beads following the manufacturer’s protocol (Beckman Coulter). RNA concentration and purity were measured using a NanoDrop Spectrophotometer (Thermo Scientific). cDNA synthesis was performed using 500 ng RNA with Maxima H Minus cDNA Synthesis Master Mix (Thermo Scientific) following the manufacturer’s protocol. The 10 µL cDNA reaction was diluted to 200 µL and 3 µL was used with iTaq Universal SYBR Green Supermix (Bio-Rad) and 670 nM of primer pairs (**Table S4**) in a 10 µL reaction for immediate use in RT-qPCR. An epMotion 5073 (Eppendorf) was used to pipette cDNA, iTaq mix, and primers into a white 96 well semi-skirted plate (MIDSCI PR-PCR2196LC-W). The plate was sealed with Avant ThermalSeal optically clear polyester RT-PCR film (MIDSCI TS-RT2-100), briefly spun down, and loaded into a LightCycler 96 Instrument (Roche). Pre-incubation at 95°C for 90 sec followed by 40 cycles of 95°C 10 sec, 57°C 10 sec, 72°C 10 sec was performed, taking fluorescence readings after the 72°C extension step was performed to obtain C_q_ values.

### NMD efficiency calculations from RT-qPCR

Once 2^-ΔΔCq^ values were obtained, we used the following to calculate NMD efficiency: 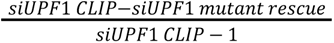 for the endogenous expression conditions and 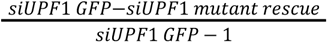 for the overexpression conditions.

## Acknowledgements

We thank members of the Hogg lab, Charles Bou-Nader, Ian Morgan, and Aparna Kishor for critical discussion and reading of this manuscript. We thank Jonathan Silver of the Neuman lab for help in deriving processivity fitting equations. We thank Anthony Armstrong of NIAID for the generous help in adapting the Rosetta-Vienna RNP ΔΔG method to the Biowulf high performance computing system and for help using Bioluminate. We thank Soumya Ranganathan and Sarah Fritz for AKTA-purified UPF1-HD protein. This work was supported by the Intramural Research Program, National Heart, Lung, and Blood Institute, National Institutes of Health, and utilized the computational resources of the NIH HPC Biowulf cluster (http://hpc.nih.gov).

## Author Contributions

J.H.C., K.C.N., and J.R.H. conceptualized the study. J.H.C. wrote the original manuscript and J.H.C., K.C.N., and J.R.H. edited it. J.H.C. and C.D.W. cloned plasmids and purified proteins. J.H.C. performed the *in silico* screen. J.H.C. acquired and analyzed all data, except for SPRNT data, which was acquired and analyzed by J.M.C. J.M.C. and J.H.G. were supported by National Human Genome Research Institute Grant R01HG005115. C.D.W. constructed initial plasmids to reduce translation of CLIP-UPF1 protein in mammalian cells.

